# Chaining for Accurate Alignment of Erroneous Long Reads to Acyclic Variation Graphs^*^

**DOI:** 10.1101/2022.01.07.475257

**Authors:** Jun Ma, Manuel Cáceres, Leena Salmela, Veli Mäkinen, Alexandru I. Tomescu

## Abstract

Aligning reads to a variation graph is a standard task in pangenomics, with downstream applications such as improving variant calling. While the `vg` toolkit (Garrison et al., *Nature Biotechnology*, 2018) is a popular aligner of short reads, `GraphAligner` (Rautiainen and Marschall, *Genome Biology*, 2020) is the state-of-the-art aligner of erroneous long reads. `GraphAligner` works by finding candidate read occurrences based on *individually* extending the best seeds of the read in the variation graph. However, a more principled approach recognized in the community is to co-linearly chain *multiple* seeds.

We present a new algorithm to co-linearly chain a set of seeds in a string labeled acyclic graph, together with the first efficient implementation of such a co-linear chaining algorithm into a new aligner of erroneous long reads to acyclic variation graphs, `GraphChainer`. Compared to `GraphAligner`, `GraphChainer` aligns 12% to 17% more reads, and 21% to 28% more total read length, on real PacBio reads from human chromosomes 1, 22 and the whole human pangenome. On both simulated and real data, `GraphChainer` aligns between 95% and 99% of all reads, and of total read length. We also show that `minigraph` (Li et al., *Genome Biology*, 2020) and `minichain` (Chandra and Jain, *RECOMB*, 2023) obtain an accuracy of less than 60% on this setting.

`GraphChainer` is freely available at https://github.com/algbio/GraphChainer. The datasets and evaluation pipeline can be reached from the previous address.

## 1 Introduction

### Motivation

Variation graphs are a popular representation of all the genomic diversity of a population [7, 12, 34]. While a single reference sequence can be represented by a single labeled path, a variation graph encodes each variant observed in the population via an “alternative” path between its start and end genomic positions in the reference labeled path. For example, the popular `vg` toolkit [15] can build a variation graph from a reference genome and a set of variants, or from a set of reference genomes, while also supporting alignment of reads to the graph.

An interesting feature of a variation graph is that it allows for recombinations of two or more variants that are not necessarily present in any of the existing reference sequences used to build the graph. Namely, the paths of a variation graph now spell novel haplotypes recombining different variants, thus improving the accuracy of downstream applications, such as variant calling [15, 47, 18, 19, 44]. In fact, it has been observed [15] that mapping reads with `vg` to a variation graph leads to more accurate results than aligning them to a single reference (since it decreases the reference bias). The alignment method of `vg` was developed for short reads [15], and while it can run also on long reads, it decreases its performance (running time, memory usage and alignment accuracy) significantly [37]. To the best of our knowledge, the only sequence-to-graph aligners tailored to long reads are `SPAligner` [11], designed for assembly graphs, `GraphAligner` [37], designed for both assembly and variation graphs, `PaSGAL` [23], designed for both short and long reads on acyclic graphs, the extension of `AStarix` for long reads [21] as well as `minigraph` [31] and the recent `minichain` [5]. `GraphAligner` is the state-of-the-art aligner for more-erroneous long reads at the whole human pangenome level^1^, allowing for faster and more accurate alignments.

### Background

At the core of any sequence-to-graph aligner is the so-called *string matching to a labeled graph* problem, asking for a path of a node-labeled graph (e.g., a variation graph) whose spelling matches a given pattern. If allowing for approximate string matching under the minimum number of edits in the sequence, then the problem can be solved in quadratic time [2], and extensions to consider affine gap costs [24] and various practical optimizations were developed later. These practical optimizations were implemented into fast exact aligners such as `PaSGAL` [23], `GraphAligner` [36, 37], `AStarix` [21, 20].

Since, conditioned on SETH, no strongly subquadratic-time algorithm for edit distance on sequences can exist [3], these sequence-to-graph alignment algorithms are worst-case optimal up to subpolynomial improvements. This lower bound holds even if requiring only *exact* matches, the graph is acyclic, i.e. a DAG [13, 16], and we allow any polynomial-time indexing of the graph [14]. Recently, the bound was shown to hold for the special case of De Bruijn graphs [17].

Given the hardness of this problem, current tools such as `vg`, `GraphAligner`, `minigraph` and `minichain` employ various heuristics and practical optimizations to approximate the sequence-to-graph alignment problem, such as partial order alignment [27], as in the case of `vg`, seed-and-extend based on minimizers [38], as in the case of `GraphAligner`, `minigraph` and `minichain`, and co-linear chaining in graphs [33] as in the case of `minigraph` and `minichain`. In the case of `GraphAligner`, the aligner finds the minimizers of the read that have occurrences in the graph, clusters and ranks these occurrences, similarly to `minimap` [29], and then tries to extend the best clusters using an optimized *banded* implementation of the bit-parallel edit distance computation for sequence-to-graph alignment [36]. This strategy was shown to be effective in mapping erroneous long reads, since minimizers between erroneous positions can still have an exact occurrence in the graph. However, this strategy can still fail. For example, the seeds may be clustered in several regions of the graph and be separated in distance. Extending from one seed (cluster) through an erroneous zone to reach the next relatively accurate region would be hard in this case. Hence, for a long read, such an aligner may find several short alignments covering different parts of the read, but not a long alignment of the entire read. Furthermore, a seed may have many false alignments in the graph that are not useful in the alignment of the long read, but which can only be discarded when looking at its position compared to other seed hits. A standard way to capture the global relation between the seed hits is through the *co-linear chaining* problem [35], originally defined for two sequences as follows. Given a set of *anchors* consisting of pairs of intervals in the two sequences (e.g., various types of matching substrings, such as minimizers), the goal is to find a *chain* of anchors such that their intervals appear in the same order in both sequences. If the goal is to find a chain of maximum coverage in one of the sequences, the problem can be solved in time *O*(*N* log *N*) [1, 42, 32], where *N* is the number of anchors. Co-linear chaining is successfully used by the popular read-to-reference sequence aligner `minimap2` [30] and also in `uLTRA` [32], an aligner of RNA-seq reads. Moreover, recent results show connections between co-linear chaining and classical distance metrics [32, 22]. The co-linear chaining problem can be naturally extended to a sequence and a labeled graph and has been previously studied for DAGs [33, 26], but now considering the anchors to be pairs of a path in the graph and a matching interval in the sequence. `GraphAligner` refers to chaining on DAGs as a principled way of chaining seeds [37, p. 16], but does not implement co-linear chaining. Also `SPAligner` appears to implement a version of co-linear chaining between a sequence and an assembly graph. It uses the BWA-MEM library [28] to compute anchors between the long read and individual edge labels of the assembly graph, and then extracts “the heaviest chain of compatible anchors using dynamic programming” [11, p.3– 4]. Their compatibility relation appears to be defined as in the co-linear chaining problem, and further requires that the distance between the anchor paths is relatively close to the distance between anchor intervals in the sequence. However, the overall running time of computing distances to check compatibility is quadratic [11, Supplemental material]. In the case of `minigraph`, the tool also computes seeds using minimizers and then applies a two-round co-linear chaining strategy to chain them. In the first round it chains seeds within a node using a `minimap2`-like chaining algorithm, while in the second round it uses the chaining scores computed previously to chain the linear chains in the graph. Finally, the recent tool `minichain` showed how to improve `minigraph`’s chaining method by following a more principled approach on their chaining algorithm.

If the co-linear chaining problem is restricted to **character**-labeled DAGs, it can be solved in time *O*(*kN* log *N* + *k* |*V*|), after a preprocessing step taking *O*(*k*(|*V*| + |*E*|) log *V*) time [33] when suffix-prefix overlaps between anchor paths are not allowed. Here, the input is a DAG *G* = (*V, E*) of *width k*, which is defined as the cardinality of a minimum-sized set of paths of the DAG covering every node at least once (a *minimum path cover*, or MPC). Thus, if *k* is constant, this algorithm matches the running time of co-linear chaining on two sequences, plus an extra term that grows linearly with the size of the graph. Even though there exists an initial implementation of these ideas tested on (small) RNA splicing graphs [26], it does not scale to large graphs, such as variation graphs, because of the *k* |*V*| term spent when aligning each read. Moreover, in the same publication, the authors show how to solve the general problem when suffix-prefix path overlaps of any length are allowed with an additional running time of *O*(*L* log^2^ |*V*|) or *O*(*L* + #*o*)^2^, by using advanced data structures (FM-index, two-dimensional range search trees, generalized suffix tree).

### Contributions

In this paper we compute for the first time the width of variation graphs of all human chromosomes of thousands of individuals, which we observe to be at most 9 (see Table 4 in the Appendix)^3^, further motivating the idea of parameterizing the running time of the co-linear chaining problem on the width of the variation graph.

**Table 1:**
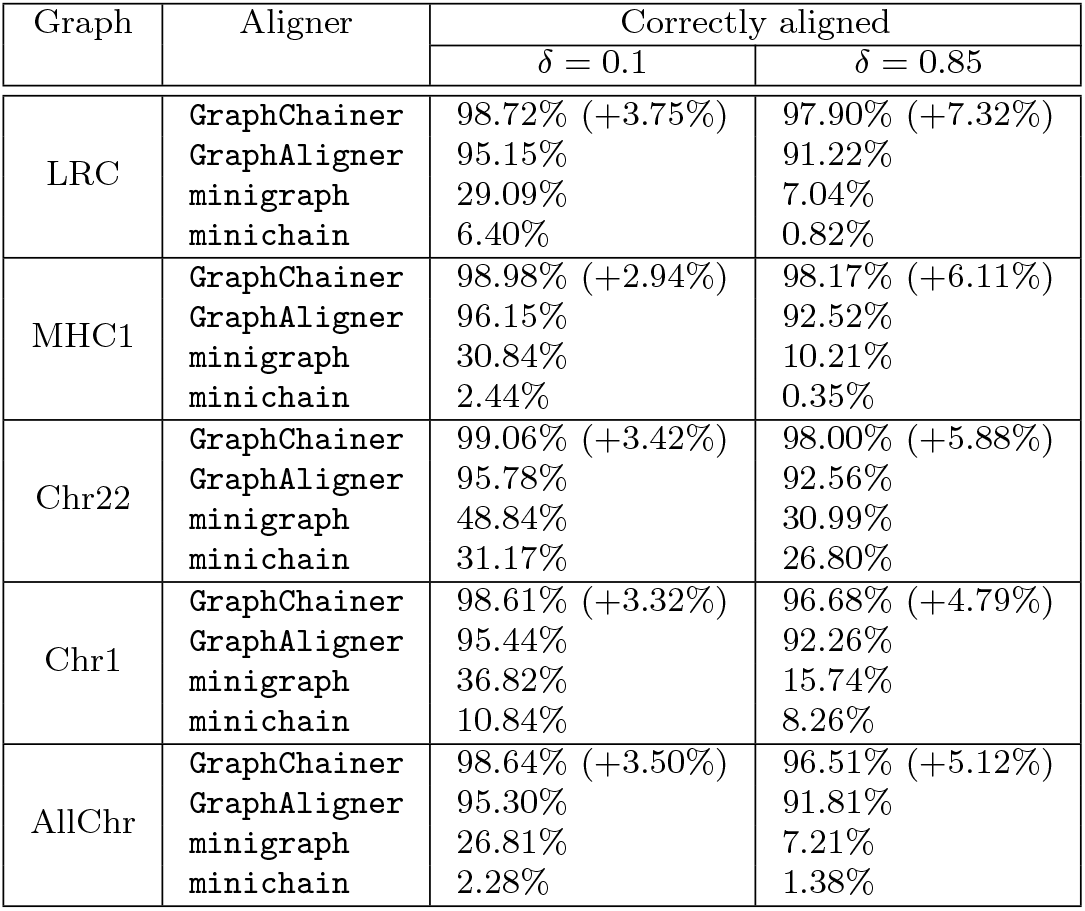
Correctly aligned reads with respect to the overlap for *δ* ∈ {0.1, 0.85} (i.e., the overlap between the reported path and the ground truth is at least 10% or 85% of the length of the ground truth sequence, respectively) for the simulated read sets with 15% error rate. Percentages in parentheses are relative improvements w.r.t. `GraphAligner`.

| Graph | Aligner | Correctly aligned |  |
| --- | --- | --- | --- |
| | | $\delta = 0.1$ | $\delta = 0.85$ |
| LRC | **GraphChainer** | 98.72% (+3.75%) | 97.90% (+7.32%) |
|  | **GraphAligner** | 95.15% | 91.22% |
|  | **minigraph** | 29.09% | 7.04% |
|  | **minichain** | 6.40% | 0.82% |
| MHC1 | **GraphChainer** | 98.98% (+2.94%) | 98.17% (+6.11%) |
|  | **GraphAligner** | 96.15% | 92.52% |
|  | **minigraph** | 30.84% | 10.21% |
|  | **minichain** | 2.44% | 0.35% |
| Chr22 | **GraphChainer** | 99.06% (+3.42%) | 98.00% (+5.88%) |
|  | **GraphAligner** | 95.78% | 92.56% |
|  | **minigraph** | 48.84% | 30.99% |
|  | **minichain** | 31.17% | 26.80% |
| Chr1 | **GraphChainer** | 98.61% (+3.32%) | 96.68% (+4.79%) |
|  | **GraphAligner** | 95.44% | 92.26% |
|  | **minigraph** | 36.82% | 15.74% |
|  | **minichain** | 10.84% | 8.26% |
| AllChr | **GraphChainer** | 98.64% (+3.50%) | 96.51% (+5.12%) |
|  | **GraphAligner** | 95.30% | 91.81% |
|  | **minigraph** | 26.81% | 7.21% |
|  | **minichain** | 2.28% | 1.38% |

**Table 2:** Correctly aligned reads with respect to the distance, and percentage of read length in correctly aligned reads, for *σ*_*read*_ = 0.3 (i.e., the edit distance between the read and the reported sequence can be up to 30% of the read length) for real PacBio read sets. Percentages in parentheses are relative improvements w.r.t. `GraphAligner`.

| Graph | Aligner | Correctly aligned | Good length |
| --- | --- | --- | --- |
| Chr22<br>(real) | **GraphChainer** | 96.48% (+17.02%) | 96.30% (+28.68%) |
|  | --step 2 | 95.31% (+15.59%) | 95.01% (+26.95%) |
|  | --step 3 | 94.34% (+14.41%) | 93.82% (+25.36%) |
|  | **GraphAligner** | 82.45% | 74.84% |
|  | minigraph | 22.38% | 33.25% |
|  | minichain | 1.52% | 1.95% |
| Chr1<br>(real) | **GraphChainer** | 95.01% (+12.74%) | 94.63% (+21.80%) |
|  | --step 2 | 93.12% (+10.50%) | 92.08% (+18.51%) |
|  | --step 3 | 91.75% (+8.87%) | 89.95% (+15.77%) |
|  | **GraphAligner** | 84.27% | 77.69% |
|  | minigraph | 25.95% | 38.40% |
|  | minichain | 0.84% | 1.02% |
| AllChr<br>(real) | **GraphChainer** | 84.02% (+16.52%) | 86.75% (+26.40%) |
|  | --step 2 | 82.73% (+14.73%) | 85.05% (+23.93%) |
|  | --step 3 | 81.69% (+13.29%) | 83.53% (+21.62%) |
|  | **GraphAligner** | 72.11% | 68.63% |
|  | minigraph | 19.42% | 28.47% |
|  | minichain | 6.37% | 7.10% |

**Table 3:**
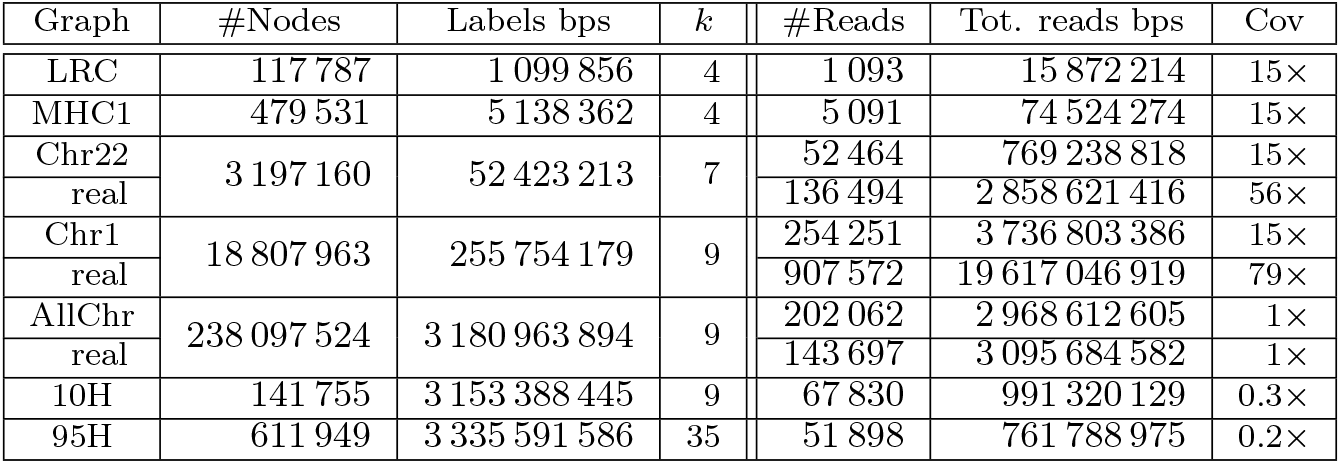
Statistics of every dataset used in our experiments. Every graph has a read set simulated on a random path of the graph. Graphs Chr22, Chr1 and AllChr were built using GRCh37 as the reference and variants from the 1000 Genomes Project, these graphs have an additional PacBio read set (rows “real”) obtained from SRA. Column *k* is the width of the graph. In the case of 10H, 95H and AllChr, *k* corresponds to the maximum width of a connected component (chromosome graph).

**Table 4:** Statistics of variation graphs of human chromosomes built with the `vg` toolkit using GRCh37 as the reference, and variants from the 1000 Genomes Project phase 3 release [6]. The number of nodes is from the compact representation where a non-branching path is merged into a single node. Each graph is a DAG, and the numbers refer only to one of the two weakly connected components (one being the reverse complement of the other). Intuitively, the width of a graph is a measure of the variability of its most complex zone, since `vg` will iteratively split those zones when building the graph. It is worth noting that these graphs are acyclic excluding structural variants.

| Chr | #Nodes | Labels bps | Width |
| --- | --- | --- | --- |
| 1 | 18 807 963 | 255 754 179 | 9 |
| 2 | 20 597 735 | 250 312 064 | 6 |
| 3 | 16 965 471 | 203 883 122 | 7 |
| 4 | 16 662 965 | 196 912 161 | 6 |
| 5 | 15 313 396 | 186 204 491 | 6 |
| 6 | 14 596 952 | 176 169 819 | 9 |
| 7 | 13 707 868 | 163 880 288 | 8 |
| 8 | 13 370 501 | 150 986 669 | 8 |
| 9 | 10 355 761 | 144 794 206 | 7 |
| 10 | 11 595 921 | 139 553 125 | 6 |
| 11 | 11 760 609 | 139 076 341 | 7 |
| 12 | 11 239 568 | 137 745 335 | 6 |
| 13 | 8 304 603 | 118 040 865 | 6 |
| 14 | 7 714 611 | 110 019 784 | 7 |
| 15 | 7 045 787 | 104 968 212 | 6 |
| 16 | 7 837 615 | 93 074 017 | 8 |
| 17 | 6 758 004 | 83 545 050 | 6 |
| 18 | 6 592 253 | 80 357 608 | 7 |
| 19 | 5 306 144 | 60 981 512 | 6 |
| 20 | 5 268 137 | 64 854 544 | 6 |
| 21 | 3 207 166 | 49 243 683 | 6 |
| 22 | 3 197 160 | 52 423 213 | 7 |
| X | 10 011 934 | 158 748 581 | 9 |
| Y | 184 003 | 59 435 025 | 2 |

We present the first algorithm solving co-linear chaining on **string**-labeled DAGs, when allowing one-node suffix-prefix overlaps between anchor paths. The reasons to consider such variation of the problem arise from practice. First, variation graphs used in applications usually compress unary paths, also known as unitigs [25], by storing the concatenation of their labels in a unique node representing them all. This not only allows to compress graph zones without variations, but also reduces the graph size, and therefore the running time of the algorithms that run on them. In Tables 3 and 4 (Appendix) we show that typical variation graphs have 10 times more nodes in their character based representation (columns #Nodes and Labels bps). Second, as discussed before, allowing suffix-prefix overlaps of arbitrary length significantly increases the running time of the algorithm, and also requires the use of advanced data structures incurring in an additional penalty in practice. However, the important special non-overlapping case in the character based representation translates into overlaps of at most one node in the corresponding string based representation (see Figure 1), which turns out to be easier to solve. Additionally, we show that allowing such overlaps suffices to solve co-linear chaining in graphs with cycles in the same running time as our solution for DAGs (Appendix).

**Figure 1:**
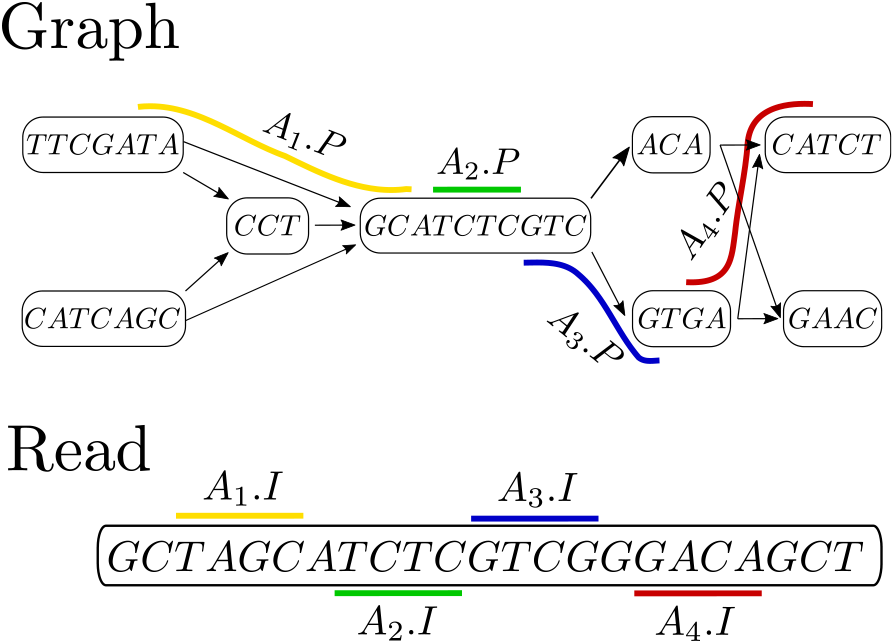
An illustrative example of Problem 2 consisting of a string labeled DAG, a read and a set of four anchors (paths in the graphs, paired with intervals in the read), shown here in different colors. The sequence 𝒞 = *A*_1_, *A*_2_, *A*_3_, *A*_4_ is a chain with *cov*(𝒞) = 16, and it is optimal since every other chain is subsequence of 𝒞. In particular, note that *A*_2_ precedes *A*_3_ because *A*_2_.*y* = 11 *< A*_2_.*x* = 12 and *A*_2_.*t* reaches *A*_2_.*t* = *A*_3_.*s*, but *A*_2_.*o*_*t*_ = 7 *< A*_3_.*o*_*s*_ = 8. If one-node suffix-prefix overlaps are not allowed, 𝒞 is not a chain and an optimal chain (either *A*_1_ or *A*_2_ followed by *A*_4_) would have coverage 8.

Our solution builds on the previous *O*(*k*(|*V*| + |*E*|) log |*V*| + *kN* log *N*) time solution [33], to efficiently consider the one-node overlapping case, but without the need of advanced data structures (Section 2.3). Moreover, we show how to divide the running time of our algorithm into *O*(*k*^3^ |*V*| + *k* |*E*|) for pre-processing the graph [4, 33], and *O*(*kN* log *kN*) for solving co-linear chaining (Appendix). That is, for constant width graphs (such as variation graphs), our solution takes linear time to preprocess the graph plus *O*(*N* log *N*) time to solve co-linear chaining, removing the previous dependency on the graph size and matching the time complexity of the problem between two sequences. Since in practice *N* is usually much smaller than the DAG size, this result allows for a more efficient implementation of the algorithm in practice.

We then implement our algorithm for DAGs into `GraphChainer`, a new sequence-to-variation-graph aligner. On both simulated and real data, we show that `GraphChainer` allows for significantly more accurate alignments of erroneous long reads than the state-of-the-art aligner `GraphAligner`. On simulated reads, we classify a read as *correctly aligned* if the reported path overlaps (100 · *δ*)% of the simulated region.

We show that `GraphChainer` aligns approximately 3% more reads than `GraphAligner` in all graphs tested and for every criterion 0 *< δ* ≤ 1. Moreover, if the criterion matches that of the average identity between the read and their ground truth, this difference increases to approximately 6% on average. On real reads, where the ground truth is not available, we classify a read as *correctly aligned* if the edit distance between the read and the reported sequence (without edits applied) is at most (100 · *σ*)% of the read length. For criterion *σ* = 0.3, `GraphChainer` aligns between 12% to 17% more real reads, of 21% to 28% more total length, on human chromosomes 1, 22 and the whole human pangenome. At the same criterion, `GraphChainer` aligns 95% of all reads or of total read length, while `GraphAligner` worsens its accuracy to less that 85% and 78%, respectively. We also run `minigraph` and `minichain` in our experimental setting of long erroneous reads and highly variable graphs encoding thousands of haplotypes, and show that these tools obtain an accuracy of less than 60% for every graph, read set and criterion evaluated.

While this increase in accuracy comes with an increase in computational resource requirements, `GraphChainer` can run in parallel and its requirements even on the whole human pangenome graph are still within the capabilities of a modern high-performance computer. We also observe that the most time-intensive part of `GraphChainer` is obtaining the anchors for the input to co-linear chaining. This is currently implemented by aligning shorter fragments of the read using `GraphAligner`, however given the modularity of this step, in the future other methods for finding anchors could be explored, such as using a fast short-read graph aligner, recent developments in MEMs [39], or even recent advances of `GraphAligner` itself.

## 2 Methods

### 2.1 Problem definition

Given a directed acyclic graph (DAG) *G* = (*V, E*), an *anchor A* is a tuple (*P* = (*s*, 舰, *t*), *I* = [*x, y*]), such that *P* is a path of *G* starting in *s* and ending in *t*, and *I* = [*x, y*] (with *x* ≤ *y* non-negative integers) is the interval of integers between *x* and *y* (both inclusive). We interpret *I* as an interval in the read matching the label of the path *P* in *G*. If *A* is an anchor, we denote by *A*.*P, A*.*s, A*.*t, A*.*I, A*.*x* and *A*.*y* its path, path starting point, path endpoint, interval, interval starting point and interval endpoint, respectively. Given a set of anchors 𝒜 = {*A*_1_, 舰, *A*_*N*_}, a *chain* 𝒞 *of anchors* (or just *chain* for short) is a sequence of anchors 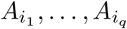 of 𝒜, such that for all *j* ∈ {1, 舰, *q* − 1}, 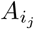 *precedes* 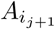, where the precedence relation corresponds to the conjunction of the precedence of anchor paths and of anchor intervals (hence *co-linear*). We will tailor the notion of precedence between paths and intervals for every version of the problem. We define the general co-linear chaining problem as follows:

#### Problem 1

(Co-linear Chaining, CLC). *Given a DAG G* = (*V, E*) *and a set* 𝒜 = {*A*_1_, 舰, *A*_*N*_} *of anchors, find an* optimal *chain* 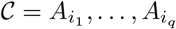 *according to some* optimization criterion.

Note that, although the final objective is to align a read to the variation graph, co-linear chaining can be defined (and solved) independent of the labels in these objects. However, in the case of string labeled graphs we will also need the exact offset *A*.*o*_*s*_ in the label of *A*.*s*, where the string represented by the anchor path starts as well as the exact offset *A*.*o*_*t*_ in the label of *A*.*t*, where the string represented by the anchor path finishes.

Co-linear chaining has been solved [33] when the optimization criterion corresponds to maximizing the coverage of the read that is 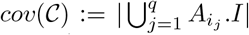, interval precedence is defined as integer inequality of interval endpoints 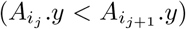, and path precedence is defined as strict reachability of path extremes (namely, 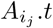 *strictly* reaches 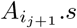, i.e., with a path of at least one edge) in time *O*(*k*(|*V*| + |*E*|) log |*V*| + *kN* log *N*). If the precedence relation is relaxed to allow suffix-prefix overlaps of paths ^4^, then their solution has an additional *O*(*L* log^2^ |*V*|) or *O*(*L* + #*o*)^5^ term in the running time. We will show how to solve co-linear chaining on string labeled DAGs when path precedence allows a one-node suffix-prefix overlap. More precisely, we solve the following problem (see Figure 1):

#### Problem 2

(One-node suffix-prefix overlap CLC). *Given a* string labeled *DAG G* = (*V, E*) *and a set* 𝒜 = {*A*_1_, 舰, *A*_*N*_} *of anchors, find a chain* 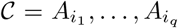*maximizing j* ∈ {,...,*q* − 1}, 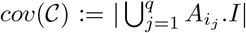, *such that for all* 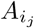 precedes 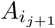, *meaning* 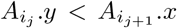 *and* 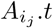 *reaches* 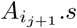, *but if* 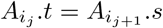 *we also require that* 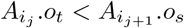.

Note that changing the strict reachability condition from the path precedence by (normal) reachability allows one-node suffix-prefix overlaps between anchor paths^6^, and no more than that (since *G* is a DAG). Also note that our definition does not allow sequence overlaps (nor in the graph nor in the sequence), which will be reflected as a simplification in our algorithm.

### 2.2 Overview of the existing solution (without suffix-prefix overlaps)

The previous solution [33] computes an MPC of *G* in time *O*(*k*(|*V* | + |*E*|) log |*V* |), which can be reduced by a recent result [4] to parameterized linear time *O*(*k*^3^|*V* | + |*E*|). Then, the anchors from A are sorted according to a topological order of its path endpoints (*A*.*t*) into *A*_1_, 舰, *A*_*N*_ . The algorithm uses a dynamic programming algorithm to compute for every *j* ∈ {1, 舰, *N*} the array:

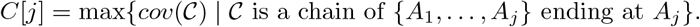

Since chains (for this version of the problem) do not have suffix-prefix overlaps, they are all subsequences of *A*_1_, 舰, *A*_*N*_, thus the optimal chain can be obtained by taking one of maximum *C*[*j*].

#### Algorithm 1

Our solution to Problem 2. For a vertex *u*, the entry `forward[u]` contains the pairs (*v, P*_*i*_), such that *u* is the last vertex, in path *P*_*i*_, that reaches *v*. These links can be pre-computed in time *O*(*k* |*E* |), and there are *O*(*k* |*V*|) of them in total [33]. Data structures 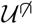 and 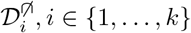, *i* {1, …, *k*} can answer `update` and `rMq` (range maximum queries).

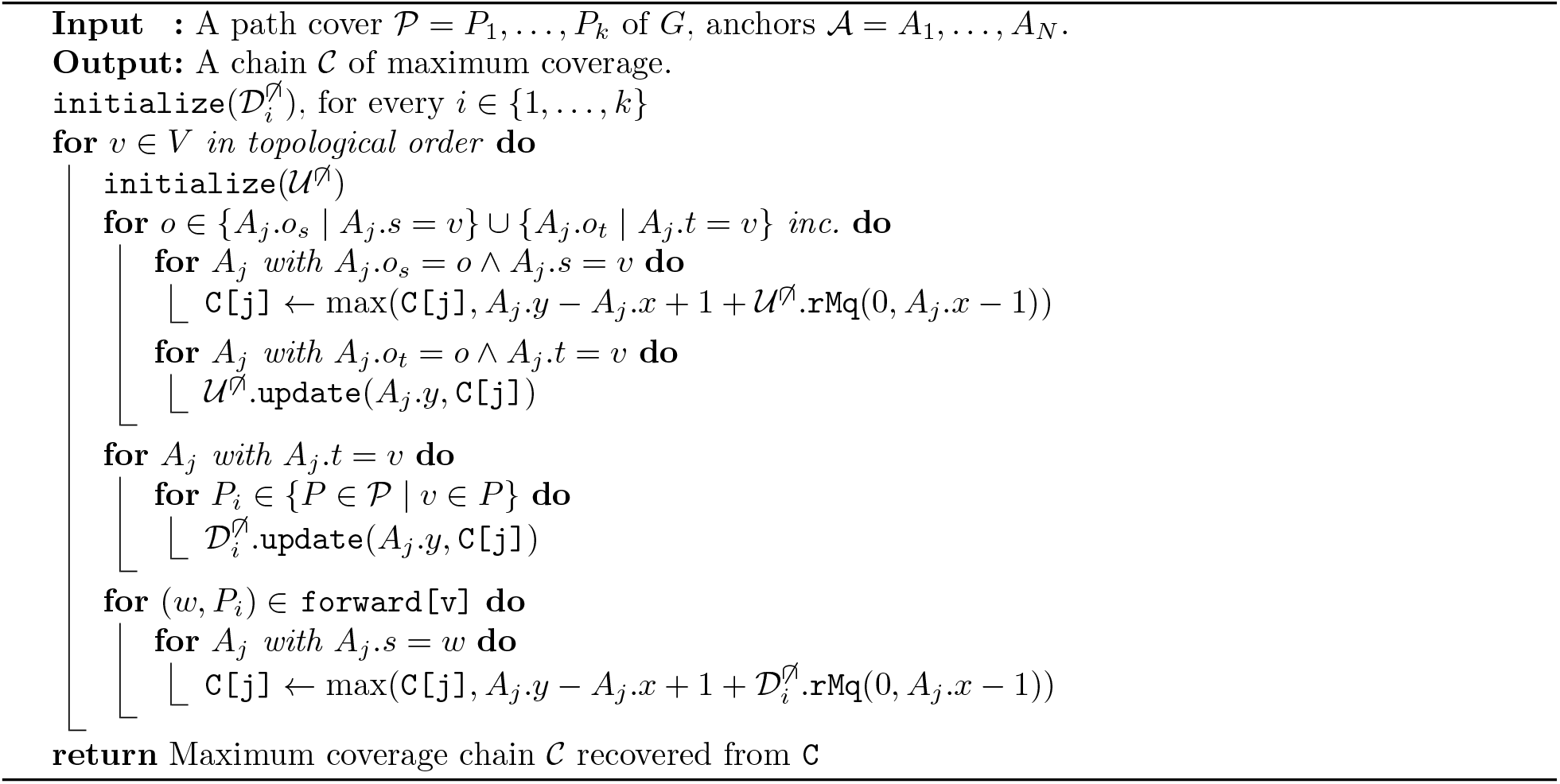

To compute *C*[*j*] efficiently, the algorithm maintains two rMq (range maximum query) data structures per path *i* in the MPC. One of the data structures, 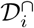, is used to compute the optimal chain whose last anchor interval overlaps the previous, and the other, 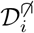, to compute the optimal chain whose last anchor interval does not overlap the previous. Since we do not consider sequence overlap, we will only use 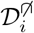 The data structure of the *i*-th path of the MPC, 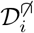, stores the information on the *C* values^7^ for all the anchors whose path endpoints belong to the *i*-th path. Since the MPC, by definition, covers all the vertices of *G*, to compute a particular *C*[*j*] it suffices to query all the data structures that contain anchors whose path endpoint reaches *A*_*j*_.*s* [33, Observation 1], which is done through the *forward propagation links*^8^.

More precisely, the vertices *v* of *G* are processed in topological order:

#### Step 1, update structures

For each anchor *A*_*j*_ whose path endpoint is *v*, and for each *i* such that *v* belongs to the *i*-th path of the MPC, the data structure of the *i*-th path is updated with the information of *C*[*j*].

#### Step 2, update *C* values

For each forward link (*u, i*) from *v* (that is, *u* is the last vertex, different from *v*, that reaches *v* in the *i*-th path), and each anchor *A*_*j*_ whose path starting point is *u*, the entry *C*[*j*] is updated with the information stored in the data structure of the *i*-th path.

Finally, since each data structure is queried and updated *O*(*kN*) times, and since these operations can be implemented in *O*(log *N*) time, the running time of the two steps for the entire computation is *O*(*kN* log *N*) plus *O*(*k*|*V* |) for scanning the forward links.

### 2.3 One-node suffix-prefix overlaps

When trying to apply the approach from Section 2.2 to solve Problem 2 we have to overcome some problems. First, it no longer holds that every possible chain is a subsequence of *A*_1_, …, *A*_*N*_ (the input 𝒜 sorted by topological order of anchor path endpoints): an anchor *A* whose path is a single vertex *v* can be preceded by another anchor *A*′ whose path ends at *v*, however *A* could appear before *A*′ in the order (for example *A*_1_ and *A*_2_ in Figure 1 could have appear in another order). To solve this issue we solve ties of path endpoints (*A*.*t*) by further comparing offsets in the string label of the path endpoint (*A*.*o*_*t*_). As such, anchors appearing after cannot precede anchors appearing before in the order, and the optimal chain can be obtained by computing the maximum *C*[*j*]. The sorting of anchors by this criterion can be done in *O*(*N* log *N*) time as it just corresponds to a sorting of the *N* anchors with a different comparison function, thus maintaining the asymptotic running time of *O*(*k*(|*V*|+ |*E*|) log |*V*| + *kN* log *N*).

A second problem is that **Step 1** assumes that *C*[*j*] contains its final value (and not an intermediate computation). Since suffix-prefix path overlaps were not allowed, this was a valid assumption in the previous case, however, because one-node suffix-prefix overlaps are now allowed, the chains whose *A*_*j*_.*P* has a suffix-prefix overlap of one node (with the previous anchor) are not considered. We fix this issue by adding a new **Step 0** before **Step 1** in each iteration. **Step 0** includes (into the computation of *C*[*j*]) those chains whose last anchors have a one-node suffix-prefix overlap, before applying **Steps 1 and 2. Step 0** uses one data structure (one in total, and is reinitialized in every **Step 0**), 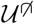, that maintains the information of the *C* values for all anchors whose paths ends at *v* (the currently processed node). In this case, the information in the data structure is not propagated through forward links (done in **Step 2** as before), but instead this information is used to update the *C* values of anchors whose paths start at *v*.

More precisely, for every anchor *A*_*j*_ whose path starts or ends at *v*, in increasing order of *A*_*j*_.*o*_*s*_ and *A*_*j*_.*o*_*t*_, respectively, we do:

#### Step 0.1, update *C* value

If the starting point of *A*_*j*_’s path is *v*, then we update *C*[*j*] with the information stored in the data structures.

#### Step 0.2, update the structure

If the endpoint of *A*_*j*_’s path is *v*, then we update the data structure with the information of *C*[*j*].

Note that in this case the update of the *C* value comes before the update of the data structures to ensure that single node anchor paths are chained correctly. Moreover, to avoid chaining two anchors *A* and *A*′ with *A*.*t* = *A*′.*s* and *A*.*o*_*t*_ = *A*′.*o*_*s*_, we process all anchors with the same offset value together, that is, we first apply **Step 0.1** to all such anchors and then **Step 0.2** to all such anchors. Algorithm 1 shows the pseudocode of the algorithm. Since every anchor requires at most two operations in the data structure the running time of **Step 0** during the whole algorithm is *O*(*N* log *N*), thus maintaining the asymptotic running time of *O*(*k*(|*V*| +| *E*|) log |*V*| + *kN* log *N*).

In the Appendix we explain how to adapt Algorithm 1 to obtain *O*(*k*^3^ |*V* |+ *k* |*E*|) time for pre-processing and *O*(*kN* log *kN*) time for solving co-linear chaining. Also in the Appendix, we show a slight variation of Problem 2 and Algorithm 1 to solve co-linear chaining on cyclic graphs.

### 2.4 Implementation

We implemented our algorithm to solve Problem 2 inside a new aligner of long reads to acyclic variation graphs, `GraphChainer`. Our C++ code is built on `GraphAligner`’s codebase. Moreover, `GraphAligner`’s alignment routine is also used as a blackbox inside our aligner as explained next (see Fig. 4 in the Appendix for an overview).

To obtain anchors, `GraphChainer` extracts *fragments* of length *ℓ* = `colinear-split-len` (default 35, experimentally set) from the long read. By default `GraphChainer` splits the input read into non-overlapping substrings of length *ℓ* each, which together cover the read. Additionally, `GraphChainer` can receive an extra parameter *s* = `sampling-step` (or just `step` for short), which samples a fragment every ⌈*s* × *ℓ* ⌉positions instead. That is, if *s >* 1, the fragments do not fully cover the input read, and if *s* ≤ 1 − 1*/ℓ*, fragments are overlapping. Having fewer fragments to align (bigger `step`) decreases the running time but also the accuracy. These fragments are then aligned with `GraphAligner` to the variation graph^9^, with default parameters; for each alignment reported by `GraphAligner`, we obtain an anchor made up of the reported path in the graph (including the offsets in the first and last nodes *A*.*o*_*s*_, *A*.*o*_*t*_), and the fragment interval in the long read^10^.

Having the input set of anchors, `GraphChainer` then solves Problem 2. The MPC index is computed with the *O*(*k*(|*V*| + |*E*|) log |*V*|) time algorithm [33] for its simplicity, where for its max-flow routine we implemented Dinic’s algorithm [10]. The `rMq` (range maximum query) data structures that our algorithm maintains per MPC path are supported by our own implementation of a treap [40].

After the co-linear chaining algorithm outputs a maximum-coverage chain 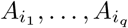, `GraphChainer` connects the anchor paths to obtain a longer path, which is then reported as the answer. More precisely, for every *j* ∈ {1, …, *q* − 1}, `GraphChainer` connects 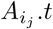 to 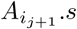 by a shortest path (in the number of nodes). Such a connecting path exists by the definition of precedence in a chain, and it can be found by running a breadth-first search (BFS) from *A*_*i*_.*t*. A more principled approach would be to connect consecutive anchors by a path minimizing edit distance, or to consider such distances as part of the optimization function of co-linear chaining. We use BFS for performance reasons.

Next, since our definition of co-linear chaining maximizes the coverage of the input read, it could happen that, in an erroneous chaining of anchors, the path reported by `GraphChainer` is much longer than the input read. To control this error, `GraphChainer` splits its solution whenever a path joining consecutive anchors is longer (label-wise) than some parameter *g* = `colinear-gap` (default 10 000, in the order of magnitude of the read length), and reports the longest path after these splits.

Finally, `GraphChainer` uses `edlib` [45] to compute an alignment between the read and the path found, and to decide if this alignment is better than the (best) alignment reported by `GraphAligner`. Therefore, `GraphChainer` can be also be viewed as a refinement of `GraphAligner`’s alignment results.

## 3 Experiments

We run several erroneous long read to variation graph alignment experiments, and compare `GraphChainer` (`v1.0.2`) against `GraphAligner` (`v1.0.13`), the state-of-the-art aligner of long reads to (pangenome level) variation graphs. We also test the performance of `minigraph` (`v0.20`) and `minichain` (`v1.0`), which also exploit co-linear chaining. We excluded `SPAligner` since this is tailored for alignments to assembly graphs. We also excluded `PaSGAL` and the recent extension of `AStarix` for long reads, since these aligners are tailored to find optimal alignments and they were three orders of magnitude slower than `GraphChainer` in our smallest dataset.

### 3.1 Datasets

#### Variation graphs

We use two (relatively) small variation graphs, two chromosome-level variation graphs, one whole human genome variation graph and other two whole human genome variation graphs used in the experiments of `minichain`. The small graphs, LRC and MHC1 [23], correspond to two of the most diverse variation regions in the human genome [9, 43]. The chromosome-level graphs, Chr22 and Chr1 (human chromosomes 22 and 1, respectively), were built with the `vg` toolkit using GRCh37 as the reference, and variants from the 1000 Genomes Project phase 3 release [6]. We use Chr22 to replicate GraphAligner’s results [37], and consider Chr1 since it is one of the most complex variation graphs of the human chromosomes (see Table 4 in the Appendix). The whole human genome graph, AllChr, corresponds to the union graph of all chromosome variation graphs, each built as described before. Finally, we use graphs 10H and 95H created for the experiments of `minichain` [5] using `minigraph` and 10 and 95 publicly available haplotype-resolved human genome assemblies.

Note that acyclic variation graphs (built as above) span all genomic positions, i.e., do not collapse repeats like, e.g., de Bruijn graphs. As such, the aligner does not perform extra steps to identify the corresponding repeat of an alignment but instead they are solved by the alignment task itself.

#### Simulated reads

For each of the previous graphs, we sample a reference sequence and use it to simulate a PacBio read dataset of 15x coverage and average error rate of 15% and 5%^11^ with the package `Badread` [48].

We use 1x coverage in the case of AllChr. To build the reference sequence we first sample a source-to-sink path from the graph by starting at a source, and repeatedly choosing an out-neighbor of the current node uniformly at random, until reaching a sink; finally, we concatenate the node labels on the path. The ambiguous characters on the simulated reference sequence are randomly replaced by one of its indicated characters. For each simulated read, we know its original location on the sampled reference sequence, which can be mapped to its ground truth location on the graph.

#### Real reads on chromosome-level graphs and whole human genome graph

For the chromosome-level graphs, we also used the whole human genome PacBio Sequel data from HG00733 (SRA accession SRX4480530)^12^. We first aligned all the reads against GRCh37 with `minimap2` and selected only the reads that are aligned to chromosome 22 and 1, respectively, with at least 70% of their length, and have no longer alignments to other chromosomes. This filtering leads to 56× and 79× coverage on Chr22 and Chr1, respectively. For AllChr we performed a uniformly random sample of all reads in the dataset to obtain 1x coverage. Table 3 (Appendix) shows more statistics of the variation graphs and read sets.

### 3.2 Evaluation metrics and experimental setup

For each read, the aligners output an alignment consisting of a path in the graph and a sequence of edits to perform on the node sequences. We call this path the *reported path* and the concatenation of the node sequences without the edits the *reported sequence*^13^. Aligners return *mapping quality scores*, however to normalize the comparison of results, we will use our own metrics to classify if an alignment was successful. On simulated data, we classify a read as *correctly aligned* if the overlap (in genomic sequences) between the reported path and the ground truth path is at least (100 ·*δ*)% of the length of the ground truth sequence, where 0 *< δ* ≤ 1 is a given threshold. As another criterion, we consider a read correctly aligned if the edit distance between the reported sequence and ground truth sequence is at most (100 · *σ*)% of the length of the ground truth sequence, where 0 *< σ* ≤ 1 is another given threshold. On real data^14^, where the ground truth is not available, we consider a read correctly aligned if the edit distance between the reported sequence and read is at most (100 *σ*)% of the read length.

Since the reads have varying sizes, we also computed the *good length*, defined as the total length of correctly aligned reads divided by the total read length, for every criterion and threshold considered.

All experiments were conducted on a server with AMD Ryzen Threadripper PRO 3975WX CPU with 32 cores and 504GB of RAM. All aligners were run using 10 threads for all datasets. Time and peak memory usage of each program were measured with the GNU `time` command. The commands used to run each tool are listed in the Appendix. Our code, datasets and pipeline can be found at https://github.com/algbio/GraphChainer.

### 3.3 Results and discussion

#### 3.3.1 Comparison with GraphAligner

##### Simulated reads with 15% error rate

For criterion *δ* = 0.85, that is matching the average identity of the simulated reads, `GraphChainer` has at least 4–5% more correctly aligned reads, and at the weaker criterion *δ* = 0.1, used in `GraphAligner`’s evaluation [37, p.3], `GraphChainer` correctly aligns 3–4% more reads, see Table 1, which is true for every criterion *δ >* 0, see Figure 2. Moreover, we observe that the accuracy of `GraphAligner` drops below 95% for *δ >* 0.6, whereas this does not happen to `GraphChainer` until *δ >* 0.98. Similar results are obtained when measuring good length and using edit distance criteria, see Tables 9, 11 and 13 and Figures 5, 6, 9, 10, 13, 14, 17 and 18 in the Appendix.

**Figure 2:**
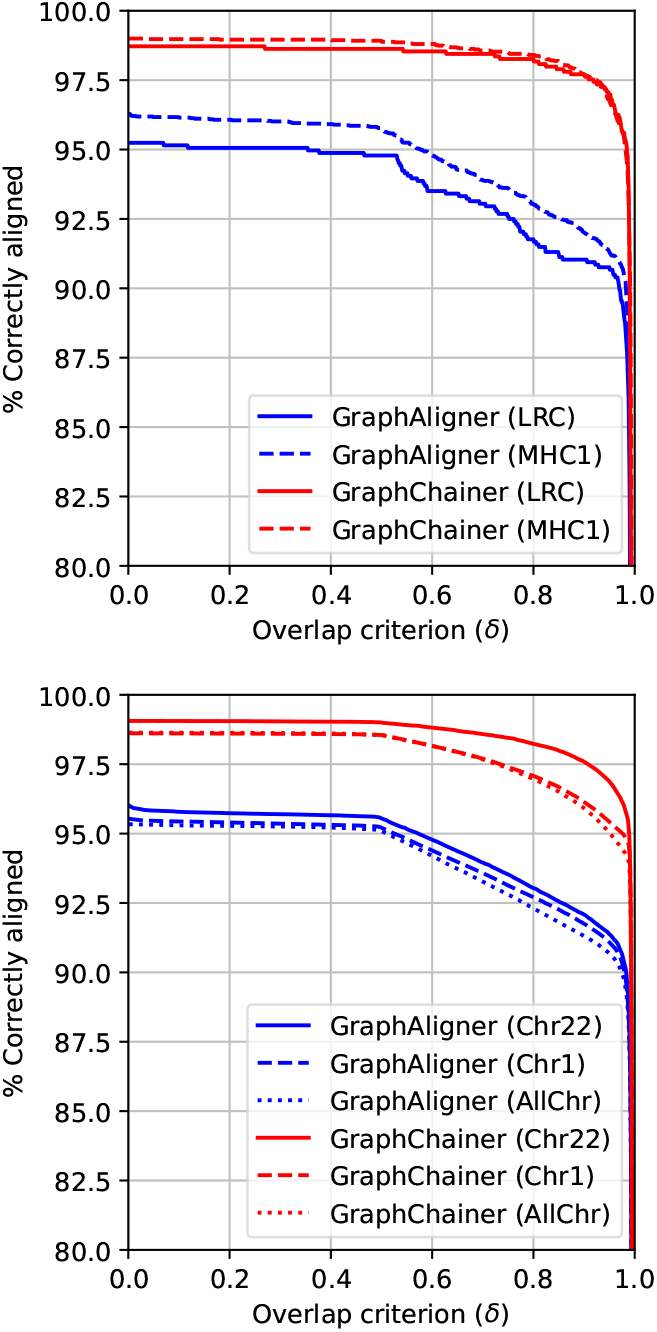
Correctly aligned reads w.r.t overlap with ground truth on the simulated read sets for LRC (top solid), MHC1 (top dashed), Chr22 (bottom solid), Chr1 (bottom dashed) and AllChr (bottom dotted). Plots for `minigraph` and `minichain` can be found in Appendix E.

**Figure 3:**
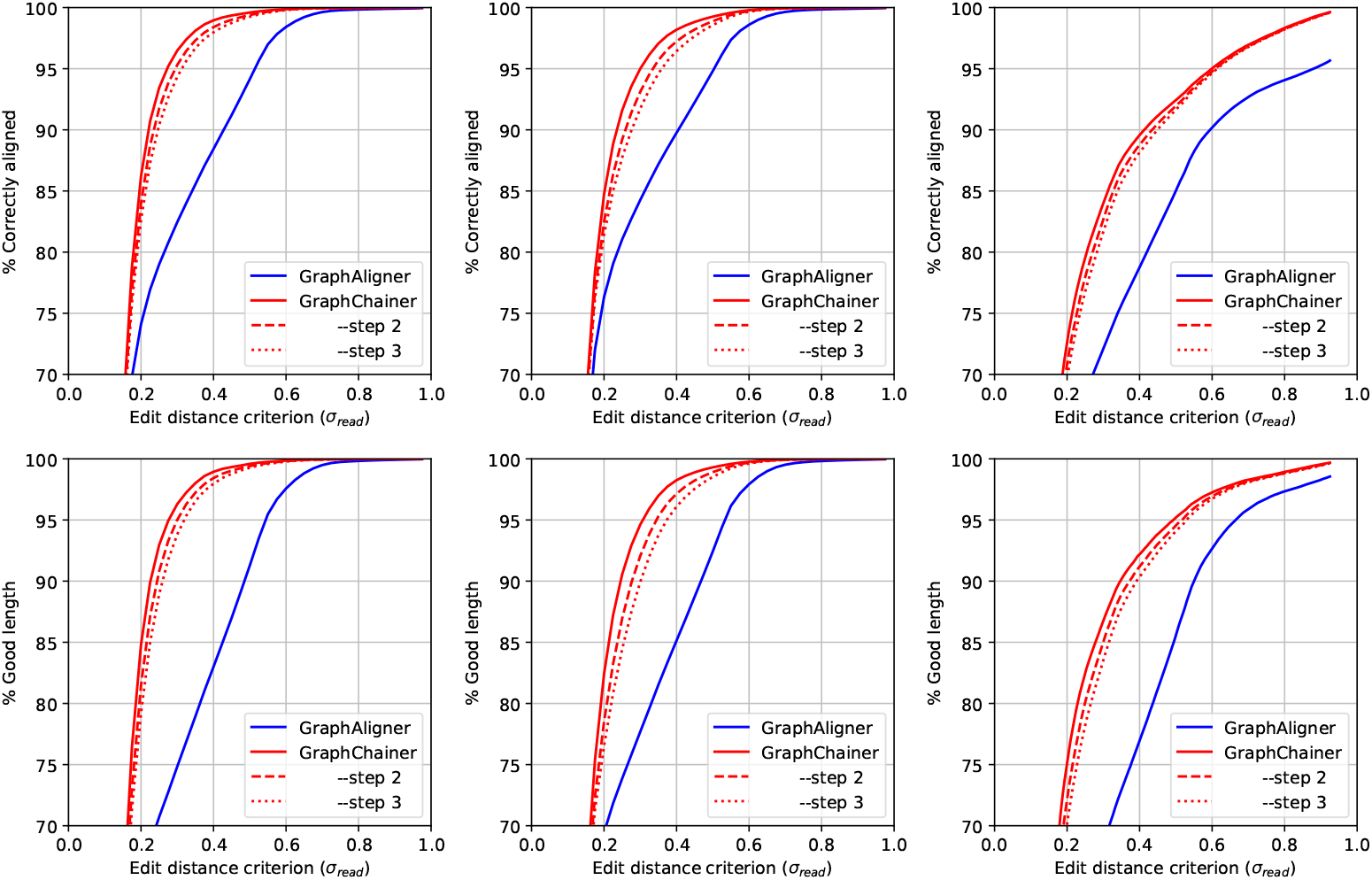
Correctly aligned reads w.r.t. the read distance (top), and read length in correctly aligned reads (bottom), on Chr22 (left), Chr1 (center) and AllChr (right), for real PacBio read sets. Plots for `minigraph` and `minichain` can be found in Appendix E

**Figure 4:**
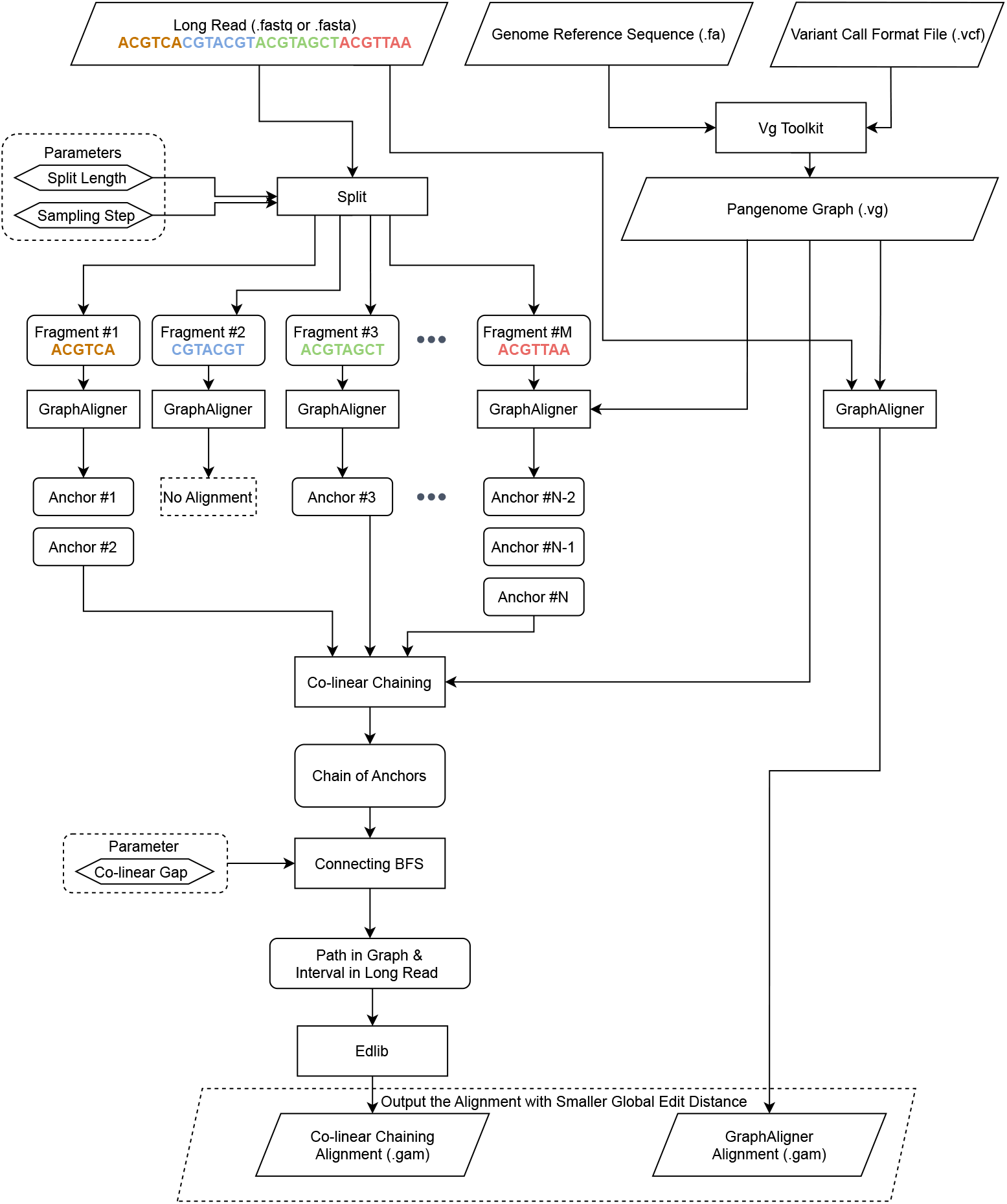
The flow diagram of `GraphChainer`: a long-read is split into fragments (of default length 35), which are aligned with `GraphAligner` (though any other method to extract anchors can be used). Anchors are created for each alignment of a fragment. The anchors are chained with our algorithm from Section 2.3. The optimal chain is split whenever the BFS shortest path between consecutive anchor paths is longer than a co-linear gap limit parameter (default 10 000), and the longest resulting path is kept. If the edit distance between the long read and this path is better than the one obtained by aligning the entire long read with `GraphAligner`, then this alignment is output; otherwise `GraphAligner`’s alignment is reported.

**Figure 5:**
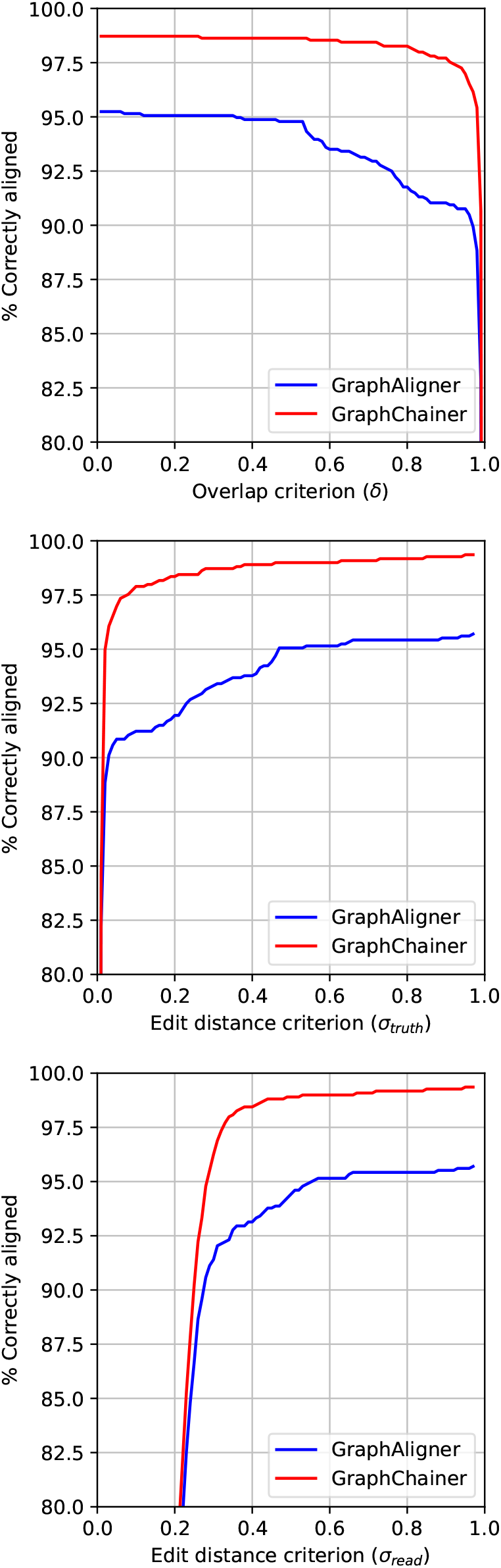
Correctly aligned reads w.r.t overlap (top), truth sequence distance (middle) (middle) and read distance (bottom) for LRC on simulated reads (only `GraphAligner` and `GraphChainer`).

**Figure 6:**
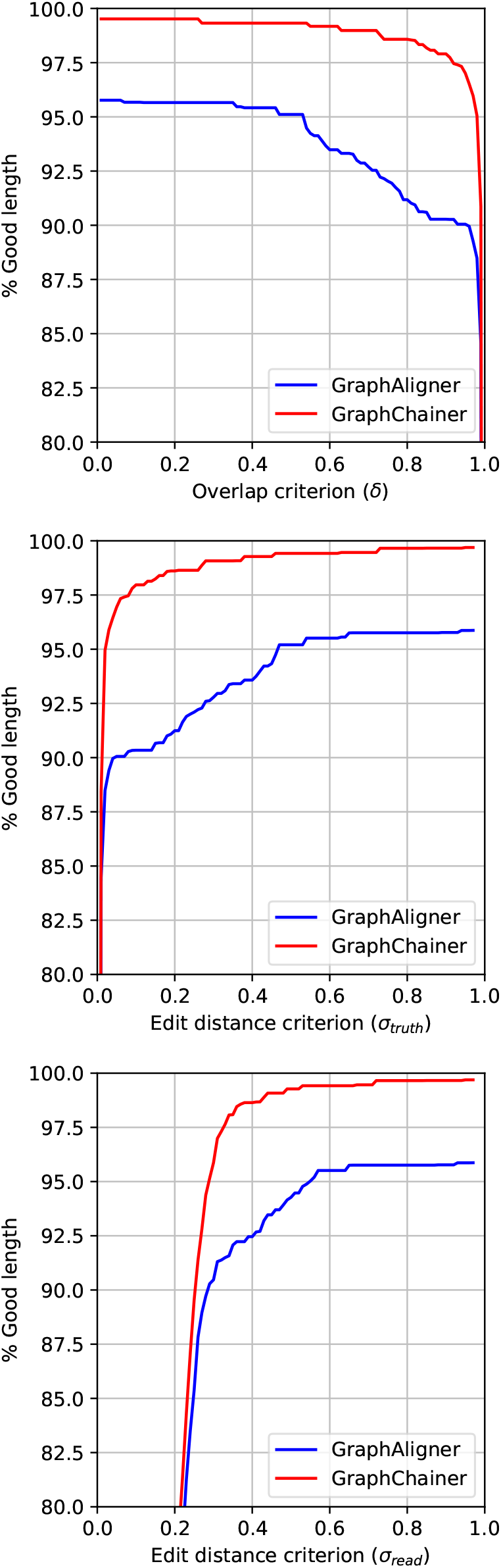
Read length in correctly aligned reads w.r.t overlap (top), truth sequence distance (middle) and read distance (bottom) for LRC on simulated reads (only `GraphAligner` and `GraphChainer`).

**Figure 7:**
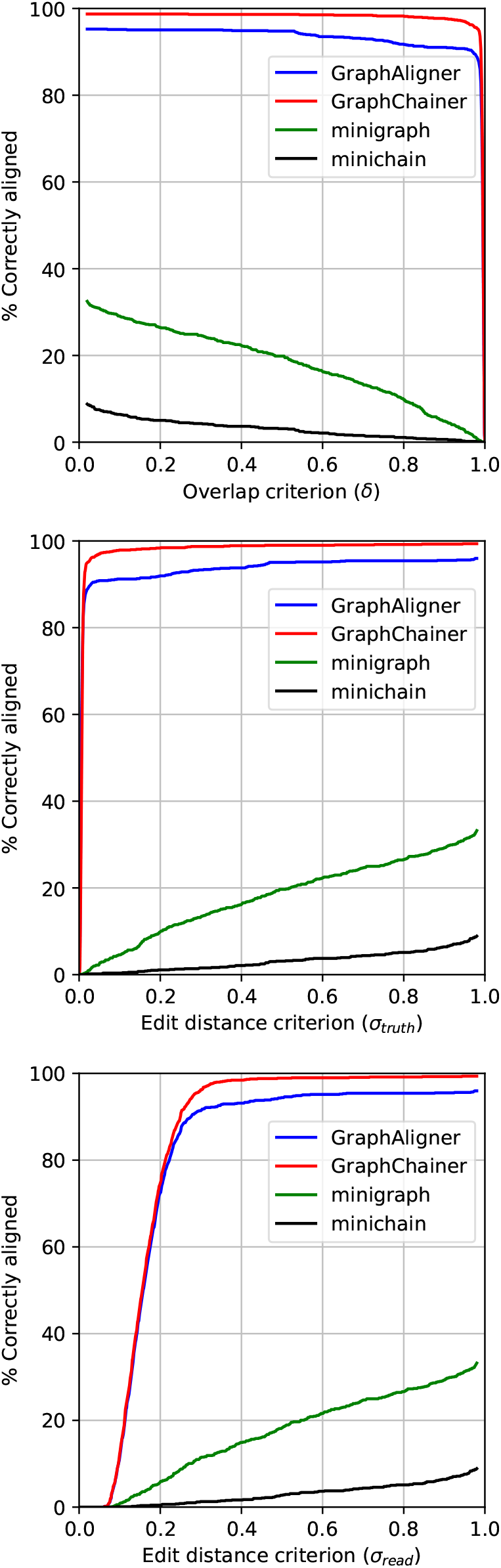
Correctly aligned reads w.r.t overlap (top), truth sequence distance (middle) (middle) and read distance (bottom) for LRC on simulated reads.

**Figure 8:**
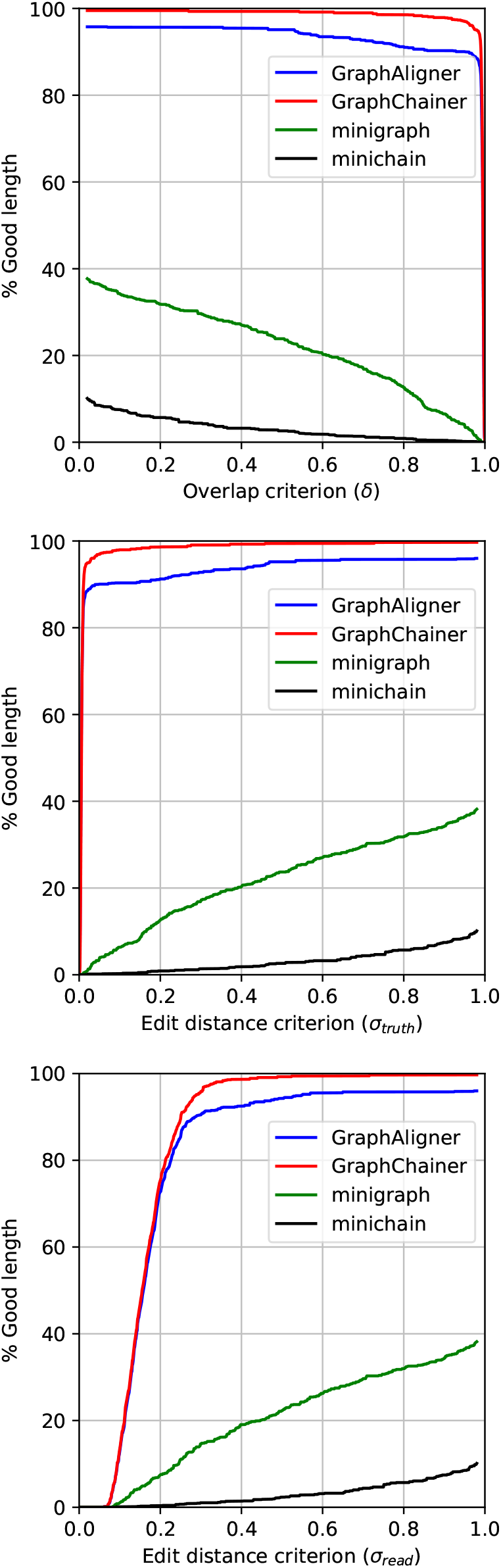
Read length in correctly aligned reads w.r.t overlap (top), truth sequence distance (middle) and read distance (bottom) for LRC on simulated reads.

**Figure 9:**
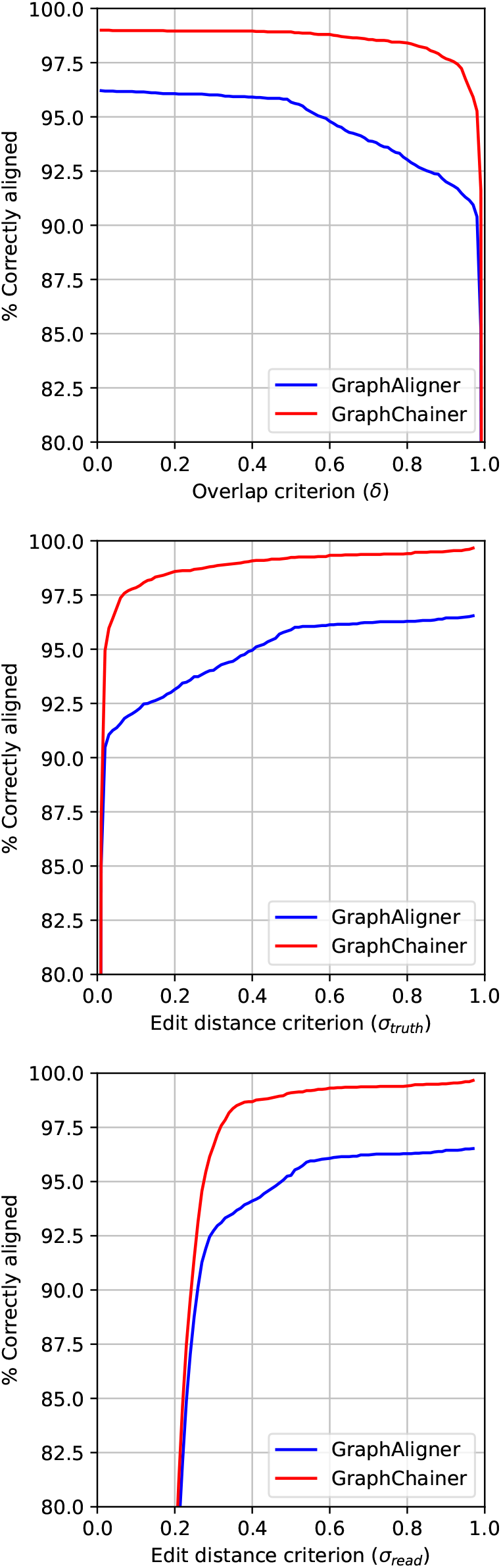
Correctly aligned reads w.r.t overlap (top), truth sequence distance (middle) and read distance (bottom) for MHC1 on simulated reads (only `GraphAligner` and `GraphChainer`).

**Figure 10:**
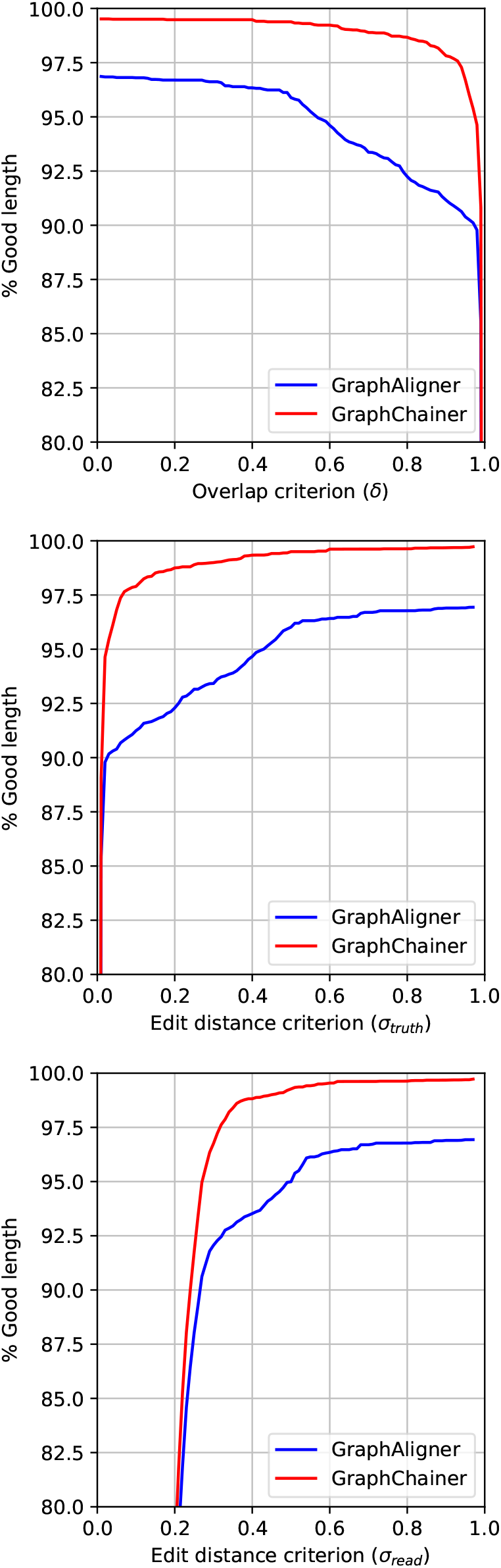
Read length in correctly aligned reads w.r.t overlap (top), truth sequence distance (middle) and read distance (bottom) for MHC1 on simulated reads (only `GraphAligner` and `GraphChainer`).

**Figure 11:**
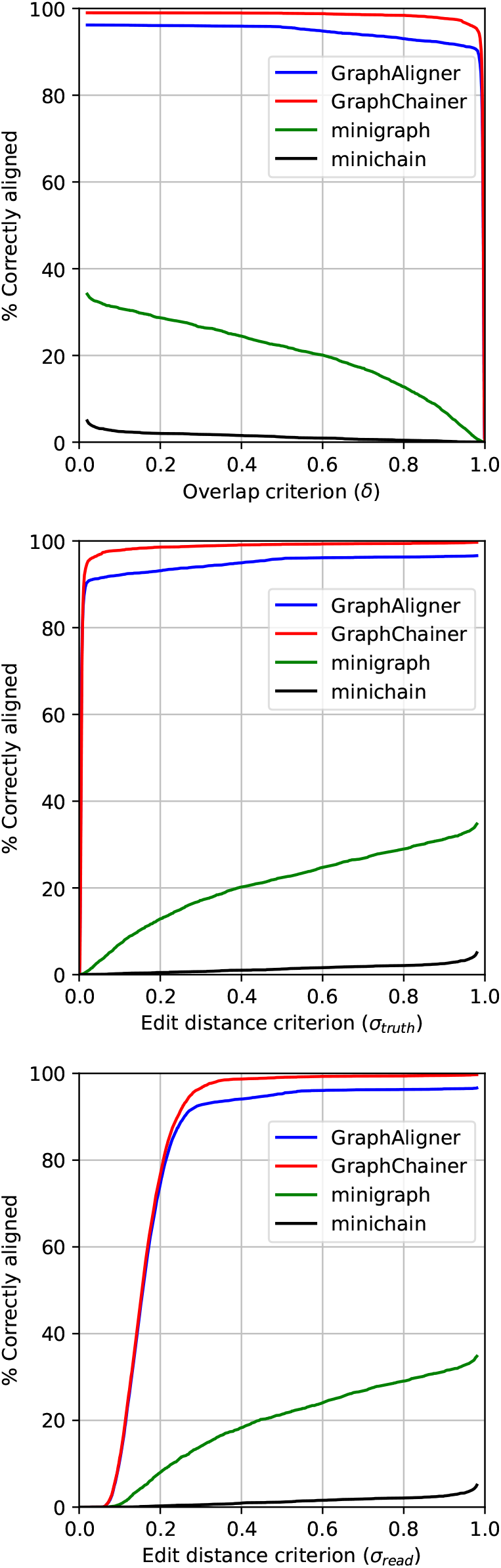
Correctly aligned reads w.r.t overlap (top), truth sequence distance (middle) and read distance (bottom) for MHC1 on simulated reads.

**Figure 12:**
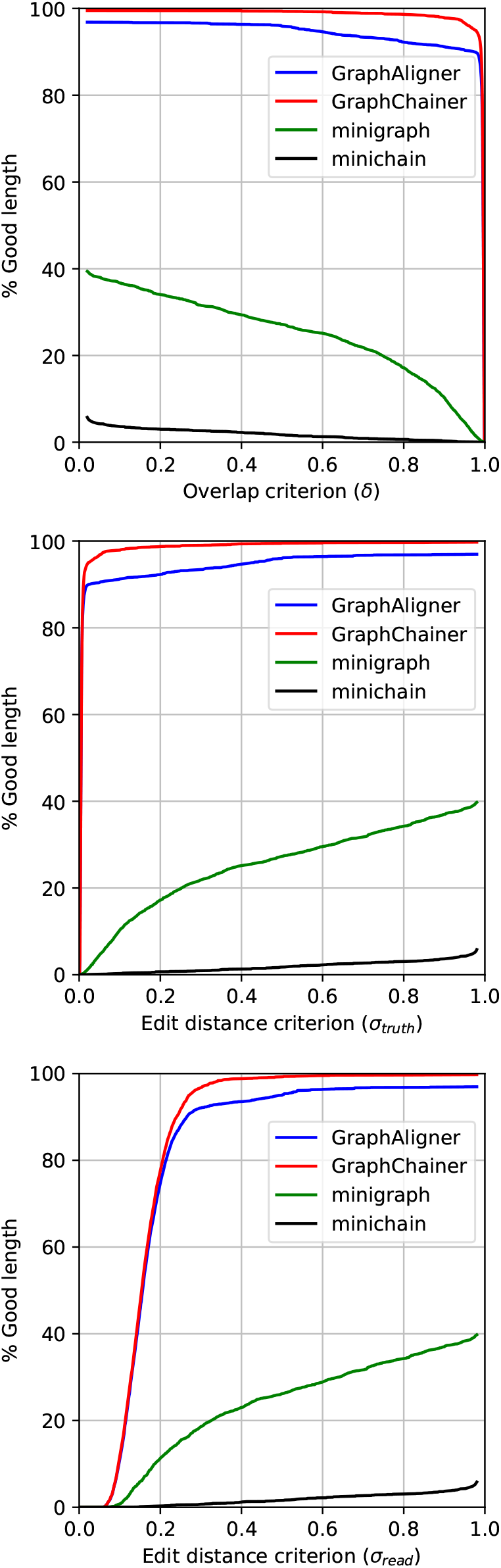
Read length in correctly aligned reads w.r.t overlap (top), truth sequence distance (middle) and read distance (bottom) for MHC1 on simulated reads.

**Figure 13:**
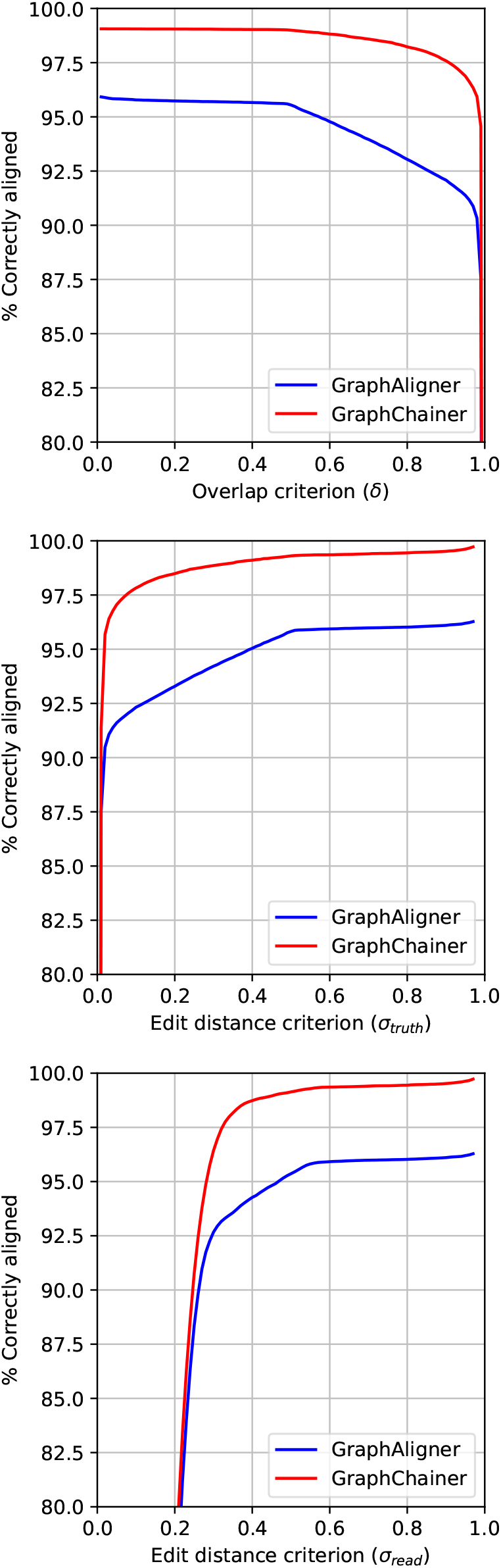
Correctly aligned reads w.r.t overlap (top), truth sequence distance (middle) and read distance (bottom) for Chr22 on simulated reads (only `GraphAligner` and `GraphChainer`).

**Figure 14:**
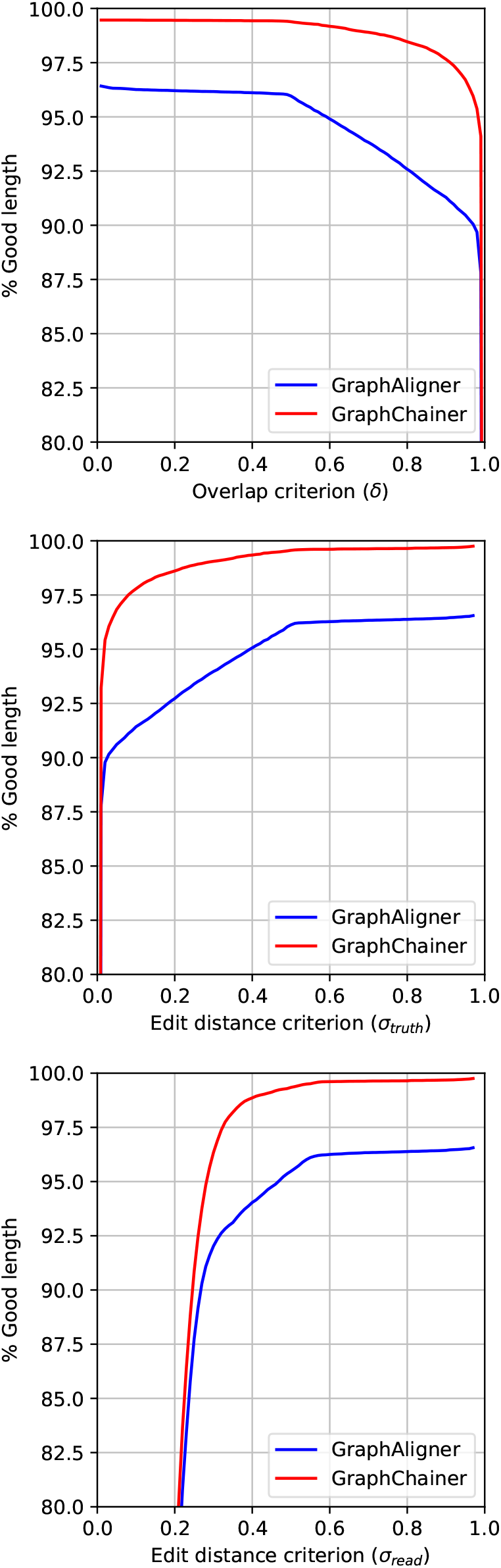
Read length in correctly aligned reads w.r.t overlap (top), truth sequence distance (middle) and read distance (bottom) for Chr22 on simulated reads (only `GraphAligner` and `GraphChainer`).

**Figure 15:**
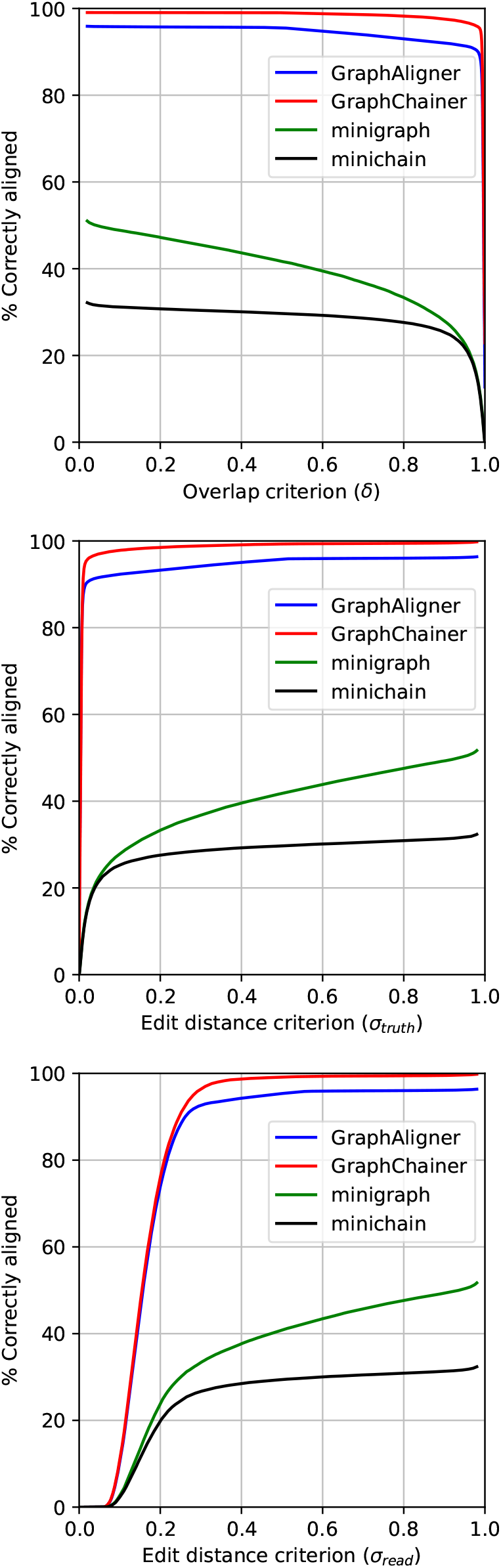
Correctly aligned reads w.r.t overlap (top), truth sequence distance (middle) and read distance (bottom) for Chr22 on simulated reads.

**Figure 16:**
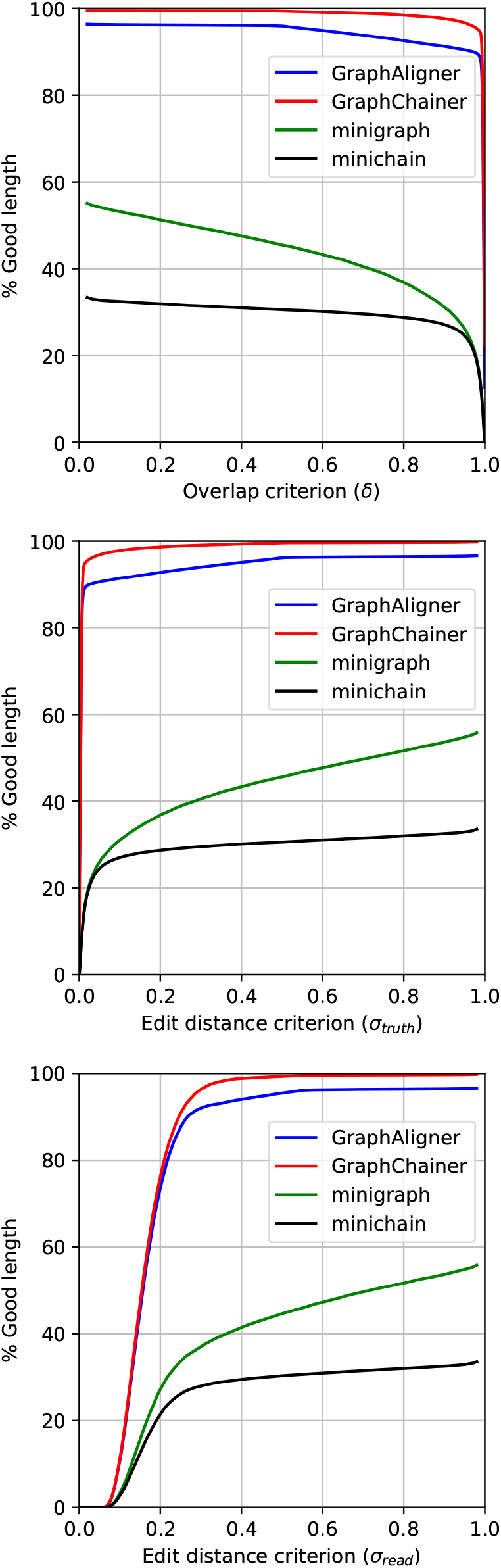
Read length in correctly aligned reads w.r.t overlap (top), truth sequence distance (middle) and read distance (bottom) for Chr22 on simulated reads.

**Figure 17:**
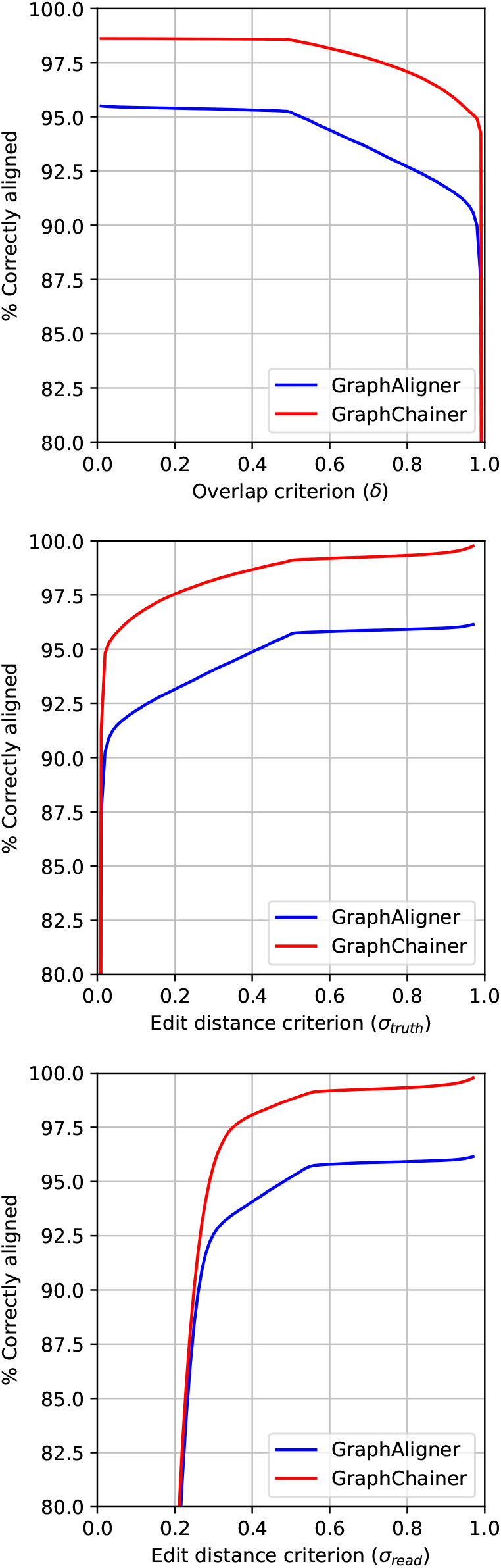
Correctly aligned reads w.r.t overlap (top), truth sequence distance (middle) and read distance (bottom) for Chr1 on simulated reads (only `GraphAligner` and `GraphChainer`).

**Figure 18:**
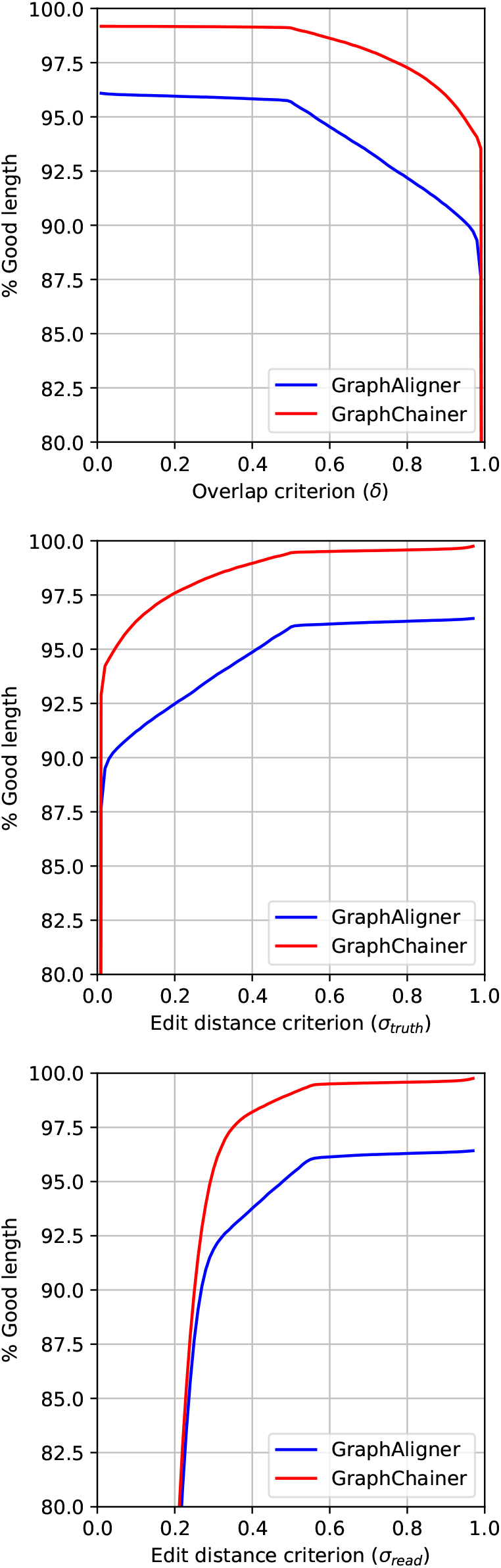
Read length in correctly aligned reads w.r.t overlap (top), truth sequence distance (middle) and read distance (bottom) for Chr1 on simulated reads (only `GraphAligner` and `GraphChainer`).

##### Real reads on chromosome and whole human genome graphs

In this case the difference between `GraphAligner` and `GraphChainer` is even clearer, see Figure 3. Since these graphs are larger, `GraphAligner`’s seeds are more likely to have false occurrences in the graph, and thus extending each seed (cluster) *individually* leads to worse alignments in more cases. As shown in Table 2, for criterion *σ* = 0.3, `GraphChainer`’s improvements in correctly aligned reads, and in total length of correctly aligned reads, are up to 17.02%, and 28.68%, respectively for Chr22. For Chr1, the improvement in the two metrics is smaller, up to 12.74%, and 21.80%, but still significant. For AllChr, the relative improvement in the metrics rises up to 16.52% and 26.40%. Both aligners decrease in accuracy when considering the *whole* human genome variation graph, but `GraphChainer` is more resilient to this change^15^. We also note that `GraphChainer` reaches an accuracy of 95% for *σ <* 0.3, whereas this happens at *σ >* 0.5 for `GraphAligner`.

#### 3.3.2 Results of minigraph and minichain

For both simulated reads with 15% error rate and real reads on all variation graphs presented in the main text `minigraph` and `minichain` align less than 60% of reads for all criteria^16^. In the case of simulated reads, at the weaker overlap criterion of *δ* = 0.1, `minigraph` correctly aligns close to 30% of reads in all graphs except Chr22 (with an accuracy of approximately 50%), and `minichain` drops to approximately 10% (except Chr22 with approximately 30% accuracy). In the case of real reads, and for criterion *σ*_*read*_ = 0.3, `minigraph` correctly aligns less than 30% of reads, but this significantly (approximately 10%) increases when considering the total read length, whereas `minichain` drops in accuracy to less than 2%.

The main reason on the low accuracy of `minigraph` and `minichain` is that these tools do not work well on the highly variable variation graphs tested (graphs with lots of small variants such as SNPs), as reported at https://github.com/lh3/minigraph#limitations, which is intensified with the high error rate of 15% on reads simulated with `Badread` [48] and the high error rate in our real PacBio reads. Results on simulated reads with lower 5% error rate^17^ show a significant improvement in these tools with `minigraph` obtaining accuracy around 90% and `minichain` around 60% for *δ* = 0.1. Moreover, at the same error rate and *δ* = 0.1, and for the (less variable) graphs 10H and 95H tested on `minichain`’s publication, `minigraph` achieves accuracy of 93.59% and `minichain` of 94.80% for 10H, and 81.94% and 83.97% for 95H, beating both `GraphAligner` and `GraphChainer`.

##### Performance

Table 6 shows the running time and peak memory of all aligners on real reads and Table 8 on simulated reads with 15% error rate (Appendix). Even though `GraphChainer` takes more time and memory resources, these are still within the capabilities of a modern high-performance computer. Running times are larger on real reads on Chr1 and AllChr due to its bigger size, but also to its larger read coverage in the case of Chr1. To explore other tradeoffs between memory and time performance and accuracy, we ran `GraphChainer` with `step` in {1, 2, 3}. We observe that the alignment accuracy remains significantly above `GraphAligner`’s, while the running time shrinks by up to half. `GraphAligner` and `minigraph` use the less memory and are the fastest. `minigraph` slightly outperforms `GraphAligner` in both metrics due to the minimizer index built by `GraphAligner`. `minichain` has a similar indexing time and peak memory as `GraphChainer`, since they are both variations of the same algorithm [33]. `minichain` has the fastest alignment time after indexing, which can be explained by the low accuracy obtained by the aligner. Table 7 in the Appendix shows that one bottleneck of `GraphChainer` is obtaining the anchors.

**Table 5:** Correctly aligned reads with respect to the overlap for *δ* ∈ {0.1, 0.85} (i.e., the overlap between the reported path and the ground truth is at least 10% or 85% of the length of the ground truth sequence, respectively) for the simulated read sets with error rate 5% as opposed to 15% shown in Table 1. Percentages in parentheses are relative improvements w.r.t. `GraphAligner`. 10H and 95H are the smallest and largest graphs used in the experiments of `minichain`, respectively.

| Graph | Aligner | Correctly aligned |  |
| --- | --- | --- | --- |
| | | $\delta = 0.1$ | $\delta = 0.85$ |
| LRC | **GraphAligner** | 98.26% | 96.24% |
|  | **GraphChainer** | 98.90% (+0.65%) | 98.72% (+2.57%) |
|  | **minigraph** | 85.60% | 67.71% |
|  | **minichain** | 45.78% | 9.72% |
| MHC1 | **GraphAligner** | 99.23% | 97.07% |
|  | **GraphChainer** | 99.64% (+0.42%) | 99.52% (+2.53%) |
|  | **minigraph** | 88.57% | 71.88% |
|  | **minichain** | 53.60% | 17.88% |
| Chr22 | **GraphAligner** | 99.12% | 97.28% |
|  | **GraphChainer** | 99.62% (+0.50%) | 99.45% (+2.24%) |
|  | **minigraph** | 91.12% | 79.88% |
|  | **minichain** | 66.49% | 41.64% |
| Chr1 | **GraphAligner** | 98.96% | 96.81% |
|  | **GraphChainer** | 99.41% (+0.46%) | 99.06% (+2.33%) |
|  | **minigraph** | 89.30% | 75.45% |
|  | **minichain** | 60.81% | 25.87% |
| AllChr | **GraphAligner** | 99.23% | 96.67% |
|  | **GraphChainer** | 99.70% (+0.47%) | 99.36% (+2.78%) |
|  | **minigraph** | 87.20% | 67.87% |
|  | **minichain** | 53.28% | 16.04% |
| 10H | **GraphAligner** | 95.28% | 93.27% |
|  | **GraphChainer** | 95.79% (+0.53%) | 95.48% (+2.38%) |
|  | **minigraph** | 93.59% | 89.72% |
|  | **minichain** | 94.80% | 92.18% |
| 95H | **GraphAligner** | 79.70% | 77.85% |
|  | **GraphChainer** | 80.48% (+0.98%) | 79.90% (+2.63%) |
|  | **minigraph** | 81.94% | 73.59% |
|  | **minichain** | 83.97% | 77.88% |

**Table 6:** Wall-clock running time (in format hh:mm:ss) and peak memory (Mem, in GBs) when aligning real PacBio read sets. Column “Map time” corresponds to the “Total time” minus indexing time.

| Graph | Aligner | Mem | Total time | Map time |
| --- | --- | --- | --- | --- |
| Chr22<br>(real) | **GraphChainer** | 8.11 | 00:27:15 | 00:26:50 |
|  | --step 2 | 7.97 | 00:21:59 | 00:21:34 |
|  | --step 3 | 8.03 | 00:20:17 | 00:19:52 |
|  | **GraphAligner** | 4.86 | 00:11:34 | 00:11:17 |
|  | **minigraph** | 5.59 | 00:41:59 | 00:41:53 |
|  | **minichain** | 8.13 | 00:04:28 | 00:03:26 |
| Chr1<br>(real) | **GraphChainer** | 38.65 | 06:39:55 | 06:37:25 |
|  | --step 2 | 38.71 | 04:18:01 | 04:15:31 |
|  | --step 3 | 38.76 | 03:31:59 | 03:29:29 |
|  | **GraphAligner** | 15.99 | 01:19:29 | 01:17:57 |
|  | **minigraph** | 12.77 | 05:04:27 | 05:03:54 |
|  | **minichain** | 31.42 | 00:57:11 | 00:48:18 |
| AllChr<br>(real) | **GraphChainer** | 419.33 | 10:09:55 | 08:12:24 |
|  | --step 2 | 419.64 | 06:22:10 | 04:24:39 |
|  | --step 3 | 421.31 | 05:08:53 | 03:11:22 |
|  | **GraphAligner** | 183.74 | 01:28:04 | 00:24:52 |
|  | **minigraph** | 117.53 | 00:53:34 | 00:45:15 |
|  | **minichain** | 394.64 | 02:27:46 | 00:12:26 |

**Table 7:** CPU time spent in the main phases of `GraphChainer` on Chr22 and Chr1 (in format hh:mm:ss) on the real PacBio reads. The column `edlib` is the time needed to run `edlib` between the read and the path found by co-linear chaining to finally report the best solution.

| Graph | step | Getting anchors | Co-linear chaining | **edlib** |
| --- | --- | --- | --- | --- |
| Chr22 | 1 | 01:19:58 | 00:25:05 | 00:46:35 |
| Chr22 | 2 | 00:39:48 | 00:10:56 | 00:48:10 |
| Chr22 | 3 | 00:26:18 | 00:06:22 | 00:49:14 |
| Chr1 | 1 | 30:29:40 | 15:12:30 | 05:39:41 |
| Chr1 | 2 | 15:15:18 | 06:38:22 | 05:50:56 |
| Chr1 | 3 | 10:08:13 | 04:03:18 | 05:54:59 |
| AllChr | 1 | 59:46:58 | 14:37:50 | 01:13:45 |
| AllChr | 2 | 29:57:32 | 06:33:21 | 01:13:50 |
| AllChr | 3 | 19:58:32 | 04:11:00 | 01:13:10 |

**Table 8:** Wall-clock running time (in format hh:mm:ss) and peak memory (Mem, in GBs) when aligning simulated reads with 15% error rate. Column “Map time” corresponds to the “Total time” minus the time taken on indexing for the respective tool.

| Graph | Aligner | Mem | Total time | Map time |
| --- | --- | --- | --- | --- |
| LRC | **GraphAligner** | 0.69 | 00:00:04 | 00:00:03 |
|  | **GraphChainer** | 0.86 | 00:00:06 | 00:00:05 |
|  | minigraph | 1.11 | 00:00:06 | 00:00:05 |
|  | minichain | 0.66 | 00:00:01 | 00:00:01 |
| MHC1 | **GraphAligner** | 1.21 | 00:00:18 | 00:00:16 |
|  | **GraphChainer** | 1.71 | 00:00:28 | 00:00:24 |
|  | minigraph | 1.75 | 00:00:30 | 00:00:29 |
|  | minichain | 1.58 | 00:00:07 | 00:00:03 |
| Chr22 | **GraphAligner** | 4.00 | 00:02:46 | 00:02:29 |
|  | **GraphChainer** | 7.38 | 00:05:25 | 00:05:00 |
|  | minigraph | 4.89 | 00:03:32 | 00:03:26 |
|  | minichain | 7.58 | 00:01:54 | 00:00:52 |
| Chr1 | **GraphAligner** | 15.75 | 00:14:36 | 00:13:04 |
|  | **GraphChainer** | 38.49 | 01:04:07 | 01:01:37 |
|  | minigraph | 12.35 | 00:21:37 | 00:21:04 |
|  | minichain | 30.69 | 00:13:57 | 00:05:04 |
| AllChr | **GraphAligner** | 183.73 | 01:28:24 | 00:25:12 |
|  | **GraphChainer** | 417.47 | 10:02:08 | 08:04:37 |
|  | minigraph | 117.21 | 00:23:14 | 00:14:55 |
|  | minichain | 394.27 | 02:23:41 | 00:08:12 |
| 10H | **GraphAligner** | 58.35 | 00:12:46 | 00:06:02 |
|  | **GraphChainer** | 87.91 | 03:02:58 | 02:54:27 |
|  | minigraph | 23.17 | 00:02:49 | 00:01:53 |
|  | minichain | 25.54 | 00:06:14 | 00:05:18 |
| 95H | **GraphAligner** | 55.85 | 00:12:14 | 00:05:08 |
|  | **GraphChainer** | 141 | 03:14:38 | 03:03:13 |
|  | minigraph | 24.74 | 00:02:59 | 00:01:58 |
|  | minichain | 24.65 | 00:12:40 | 00:11:34 |

## 4 Conclusions

The pangenomic era has given rise to several methods, including the `vg` toolkit [15], for accurately and efficiently aligning short reads to variation graphs. However, these tools fail to scale when considering the more-erroneous long reads and much less work has been published around this problem. `GraphAligner` [37] is the state-of-the-art for aligning long reads to (whole human genome) variation graphs. Here, we have presented the first efficient implementation of co-linear chaining on a string labeled DAG when allowing one-node suffix-prefix overlaps. We showed that our new method, `GraphChainer`, significantly improves the alignments of `GraphAligner`, on real PacBio reads and whole human genome variation graphs. We showed that co-linear chaining, successfully used by sequence-to-sequence aligners [30], is a useful technique also in the context of variation graphs.

One of the main drawbacks of our formulation of co-linear chaining is that the optimization criterion used by `GraphChainer`, the coverage of the read, is blind to gaps between anchor paths in the variation graph. Considering such gaps in both objects has resulted in interesting connections between co-linear chaining and classical distance metrics in the linear case [32, 22]. The recent works of `minigraph` and `minichain` have considered such gaps, showing an increase in accuracy but still relying on heuristics to approximate the optimum value of the chaining.

## A Statistics on variation graphs

## B Co-linear chaining in time *O*(*kN* log *kN*)

As we discussed, the pseudocode of Algorithm 1 takes *O*(*kN*) updates and queries to the data structures, which can be answered in *O*(log *N*) time each, thus adding up to *O*(*kN* log *N*) time in total. However, the for loops iterate over all the vertices of the graph, and over all the forward links, adding an *O*(*k*|*V*|) additional time to the chaining process.

Having such *O*(*k*|*V*|) time while chaining does not change the *O*(*k*(|*V*|+ |*E*|) log |*V*| + *kN* log *N*) asymptotic running time of our algorithm, but it affects the practical performance of chaining. As such, we decided to divide the algorithm into pre-processing the DAG, that we solve in *O*(*k*^3^|*V*| + *k*|*E*|) time by using a recent result to compute a minimum path cover of a DAG [4], and chaining that we solve in time *O*(*kN* log *kN*)^18^. To remove the chaining’s dependency on the graph size we get rid of the for loops of Algorithm 1 by instead processing the anchors (plus the required objects) in an order that simulates the for loop order of Algorithm 1. In the case of **Step 1**, this order is simulated by sorting the anchors by topological order of the path endpoints^19^, which can be done in *O*(*N* log *N*) time (by previously computing the topological ordering on pre-processing time). For **Step 0**, we also require to sort the anchors by the topological order of the path starting points, as well as interleaving the computation of **Step 0.1** and **Step 0.2**. For interleaving the computation of **Step 0.1** and **Step 0.2** (as well as the computation of **Step 0, Step 1** and **Step 2**), we partition the corresponding sorted arrays, such that each part corresponds to a different vertex, which can be done in extra *O*(*N*) time (*O*(*kN*) time in the case of **Step 2**). To interleave the computation of the different steps, we process the sorted arrays by parts according to the topological ordering of the corresponding paths. Finally, to simulate the for loop of **Step 2**, we sort the tuples (*s, P*_*i*_, *A*_*j*_) (such that (*A*_*j*_.*s, P*_*i*_) ∈ `forward[s]`) by topological order of *s*. Since there can be *O*(*kN*) such tuples, this takes *O*(*kN* log *kN*) time.

Algorithm 2 shows an alternative implementation of these ideas, which is simpler to write. Iterations of every step are encoded in 5-tuples, which are sorted to simulate the processing order of Algorithm 1. Note that |*S*| = *O*(*kN*), thus the sorting runs in time *O*(*kN* log *kN*), which dominates the running time of the algorithm. The first coordinate of the tuple is a vertex, which simulates the outer for-loop of Algorithm 1. The second coordinate is *A*_*j*_.*o*_*s*_ and *A*_*j*_.*o*_*t*_ for **Step 0.1** and **Step 0.2** and ∞ for **Step 1** and **Step 2** to simulate the for-loop of **Step 0**. The third coordinate corresponds to the `step` number and sets the relative order between the different steps. Finally, the last two coordinates are present to recover the corresponding anchor and path to process.

## C CLC in general graphs

We will show that a slight modification of our solution for Problem 2 allows us to efficiently solve an analogous definition of co-linear chaining for cyclic graphs. We will give a formal definition of this problem, show and prove how to solve it, and discuss the limitations of this approach.

In this (more general) case we allow *G* to contain directed cycles. We also allow the anchor paths to repeat vertices, and thus we change the notation *A*.*P* by *A*.*W*, where *A*.*W* is the anchor *walk* of *A*. We maintain the optimization criterion, but slightly change the definition of precedence between anchor paths to handle cyclic graphs.

### Problem 3

(CLC in general graphs). *Given a* string labeled *graph G* = (*V, E*) *and a set* 𝒜 = {*A*_1_, 舰, *A*_*N*_} *of anchors, find a chain* 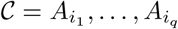 *maximizing* 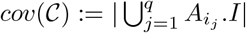, *such that for all j* ∈ {1, 舰, *q* −1}, 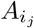 precedes 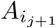, *meaning* 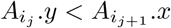 *and* 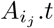 *reaches* 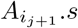, *but if* 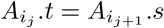 *and* 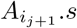 *does not strictly reaches itself we also require that* 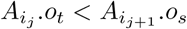.

### Algorithm 2

*O*(*kN* log *kN*) time version of Algorithm 1. A topological order on the vertices of *G* is assumed. In line 5, the 5-tuples used by the algorithm are sorted lexicographically, that is, sorting according to the first coordinate (in this case by topological order) and in case of ties sorting according to the next coordinates.

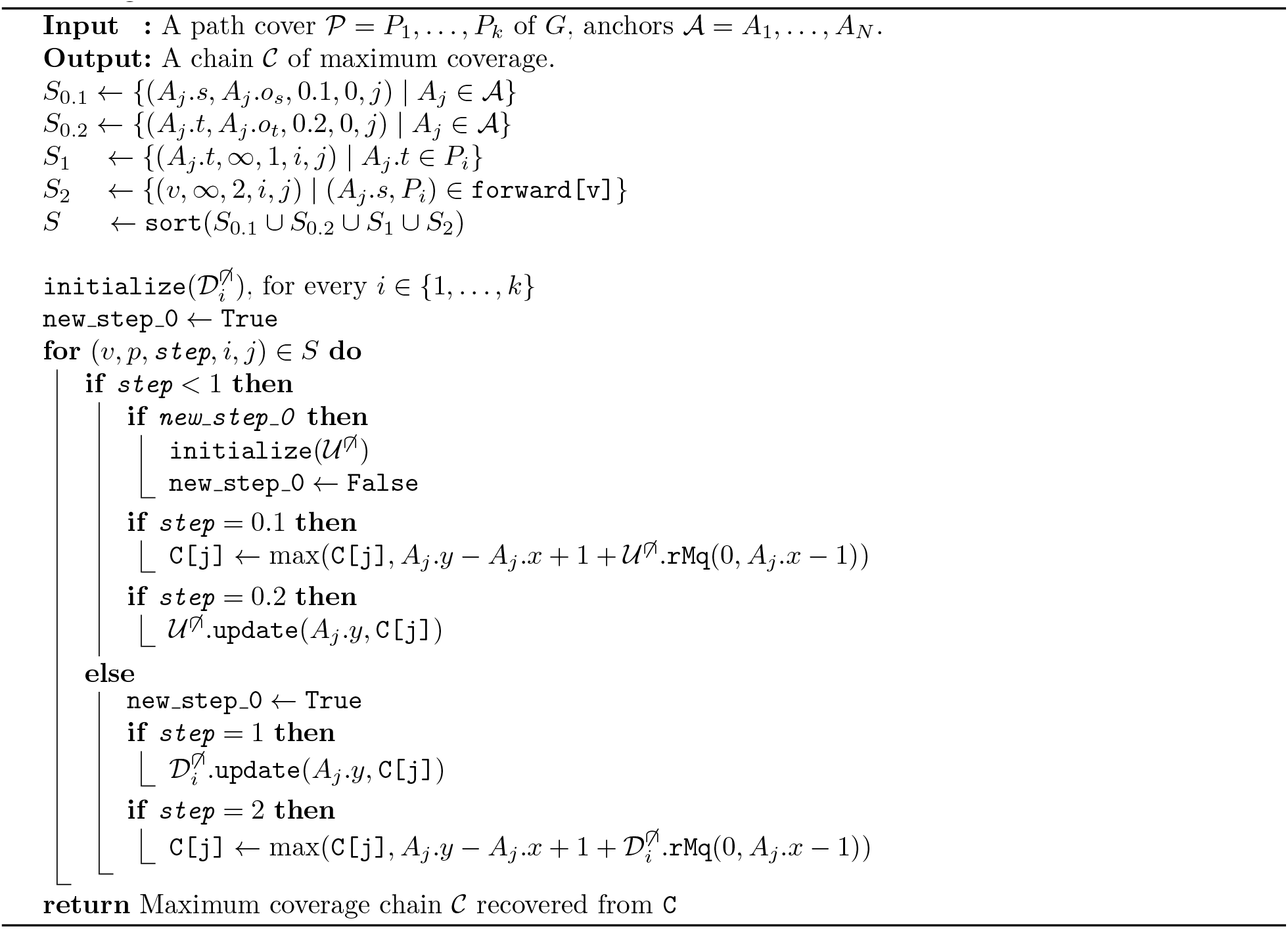

Note that the change in path precedence allows to connect paths with one-node suffix prefix overlap even when their respective offsets are inverted, but only in the case when the overlap can (strictly) reach itself. Also note that Problem 3 is a generalization of Problem 2, since in the DAG case whenever 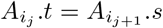 it follows that 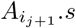 does not strictly reaches itself (otherwise we would obtain a cycle).

We solve Problem 3 by using a slight modification of our algorithm for Problem 2 in the condensation of *G*. The *condensation* = 𝒢 (𝒱, ℰ) of a graph G = (*V, E*) is the DAG of its strongly connected components, which can be computed in *O*(|*V* |+ |*E*|) time [46, 8, 41]. More precisely, 𝒱= {*S* | *S* is a strongly connected component of *G*}, and ℰ = {(*S, S*′) | ∃*u* ∈ *S, v* ∈ *S*′, (*u, v*) ∈ *E*} . Since the strongly connected components of a graph form a partition of the vertices, every vertex of *G* can be mapped to its corresponding vertex (component) in 𝒢. As such, a walk in *G* can be mapped to its corresponding path in 𝒢 (the ordered sequence of components it visits in the walk). Therefore, any path cover of *G* can be mapped into a path cover of 𝒢, thus width(*G*) = *k* ≥ *k*′ = width(𝒢). To finish our reduction we map the input anchors 𝒜 to 𝒜′ by mapping the corresponding anchor walks to paths in 𝒢. Let us call this anchor transformation *f*_𝒢_, such that, *f*_𝒢_ (*A*) = (*A*.*I, P*), where *P* is the corresponding path of *A*.*W* in 𝒢, and thus 𝒜′ = *f*_𝒢_ (𝒜). Note that this transformation is incomplete as it does not specify the offsets *f*_𝒢_ (*A*).*o*_*s*_, *f*_𝒢_ (*A*).*o*_*t*_.

If *f*_𝒢_ (*A*).*s* (resp. *f*_𝒢_ (*A*).*t*) only contains *A*.*s* (resp. *A*.*t*) then it makes sense to use *A*.*o*_*s*_ (*A*.*o*_*t*_). However, if *f*_𝒢_ (*A*).*s* (*f*_𝒢_ (*A*).*t*) contains other vertices then *f*_𝒢_ (*A*).*o*_*s*_ (*f*_𝒢_ (*A*).*o*_*t*_) is undefined since *f*_𝒢_ (*A*).*s* (*f*_𝒢_ (*A*).*t*) is not a linear structure anymore, but a subgraph. The following lemma proves that in this case it is not necessary to compute *f*_𝒢_ (*A*).*o*_*s*_ (*f*_𝒢_ (*A*).*o*_*t*_).

### Algorithm 3

Our solution to Problem 3. For a strongly connected component *u*, the entry `forward[u]` contains the pairs (*v, P*_*i*_), such that *u* is the last strongly connected component, in path *P*_*i*_, that reaches *v*. These links can be pre-computed in time *O*(*k*′|*E*|) on the condensation 𝒢 of *G*, and there are *O*(*k*′|*V* |) of them in total [33]. Data structures 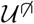 and 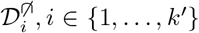, *i* ∈ {1, 舰, *k*′} can answer `update` and `rMq`. We assume that the condensation 𝒢 = (𝒱, ℰ) of *G* is already computed, as well as the mapping of anchors *f*_𝒢_.

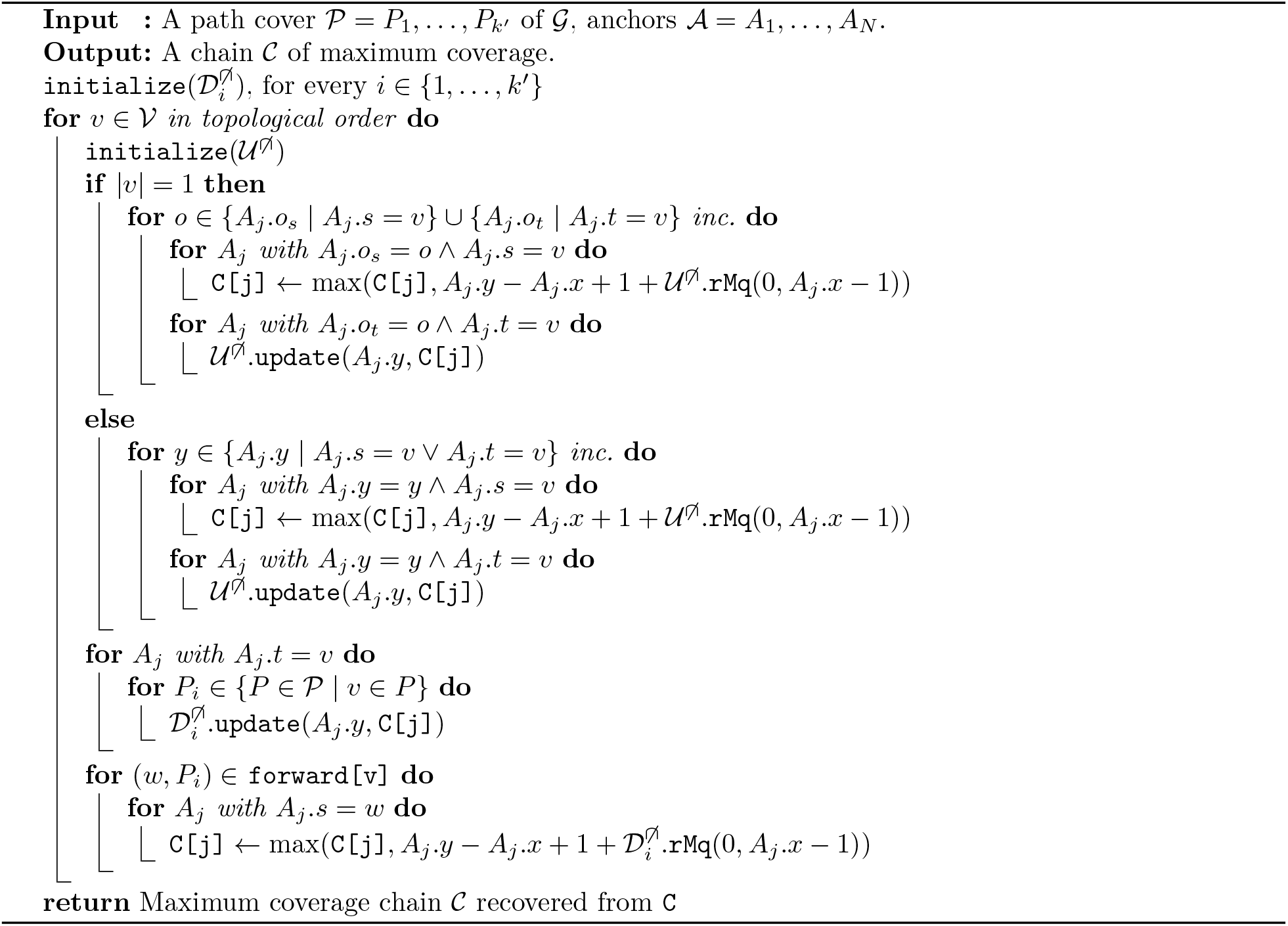

### Lemma 1.

*Following the precedence definition of Problem 3. If* |*f* _𝒢_ (*A*).*t*| *>* 1 *or* |*f* _𝒢_ (*A*′).*s*| *>* 1, *then A precedes A*′ *if and only if A*.*y < A*′.*y and A*.*t reaches A*′.*s*.

*Proof*. The forward direction of the equivalence follows by the precedence definition. The backwards direction also follows by definition if *A*.*t* ≠ *A*′.*s*. Otherwise if *A*.*t* = *A*′.*s*, note that *A*′.*s* strictly reaches itself, since |*f*_𝒢_ (*A*).*t*| = |*f*_𝒢_ (*A*′).*s*| *>* 1.

Therefore, we will proceed as in Algorithm 1 when |*f*_𝒢_(*A*).*s*| = 1 or |*f*_𝒢_(*A*).*t*| = 1, and otherwise we will run **Step 0** by increasing value of interval endpoint. Algorithm 3 shows the pseudocode for our solution, which runs in time *O*(|*V* | + |*E*| + *N* + *k*′(|𝒱| + |ℰ|) log |𝒱| + *k*′*N* log *N*) = *O*(*k*(|*V* | + |*E*|) log |*V* | + *kN* log *N*).

## D Commands for running the tools

Our code, datasets and pipeline can be found at: https://github.com/algbio/GraphChainer

Read simulation with BadReads:

~~~
badread simulate --seed {seed} --reference {Ref}
--quantity 15x --length 15000,10000
--error_model pacbio2016 --identity 85,95,5
~~~

Variation graph construction with `vg` toolkit (same parameters as those used for `GraphAligner`’s experiments [37, p.24]):

~~~
vg construct -t 30 -a -r {ref} -v {vcf} -R {chr}
-p -m 3000000
~~~

AllChr was built by taking the union of the 24 chromosome graphs.

Read alignment (same parameters for `GraphAligner` as used on its variation graph experiments [37, p.24], default parameters for `PaSGAL` [23], same parameters for the extension of `AStarix` on its experiments for long reads [21, p.13]). For `minigraph` and `minichain` we used the default parameters with the recommended option `c`:

~~~
GraphAligner -t {Threads} -x vg -f {Reads} -g {Graph}
-a {long_gam}
GraphChainer -t {Threads} -f {Reads} -g {Graph}
-a {clc_gam}
minigraph -t {Threads} -c {Graph} {Reads}
> {minigraph_gaf}
minichain -t {Threads} -c {Graph} {Reads}
> {minichain_gaf}
PaSGAL -m vg -r {Graph} -q {Reads} -t {Threads}
-o {output_file}
astarix align-optimal -a astar-seeds -g {Graph}
-q {Reads} --fixed_trie_depth 1 --seeds_len 150
-D 14 -G 1 -S 1
~~~

## E Further experimental results

**Figure 19:**
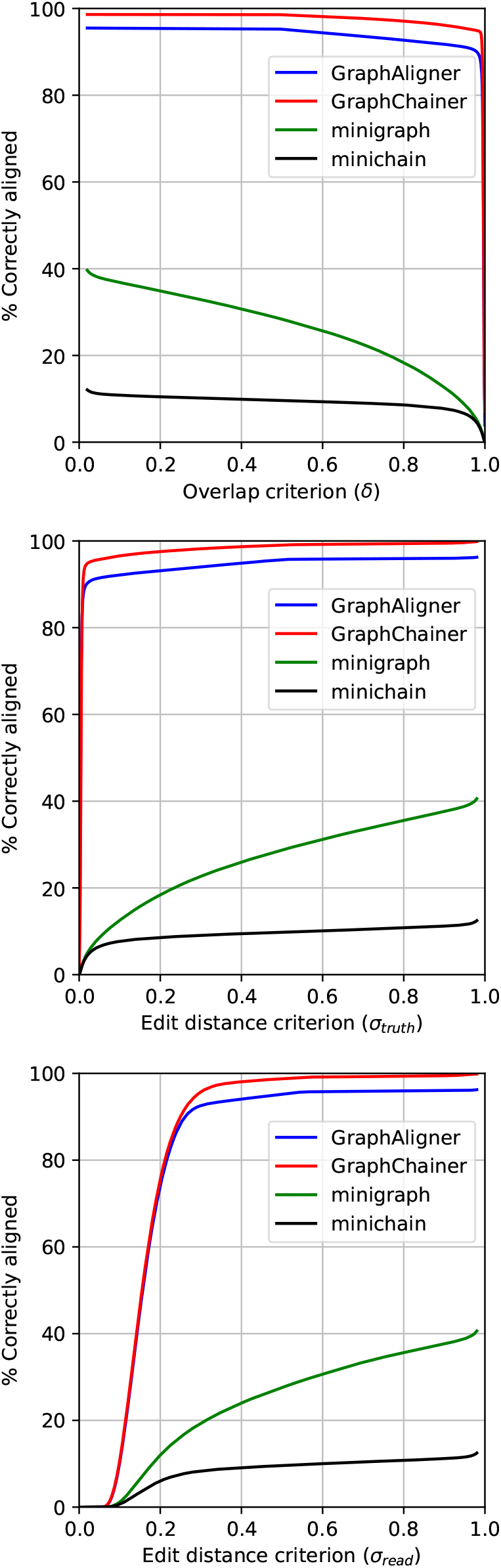
Correctly aligned reads w.r.t overlap (top), truth sequence distance (middle) and read distance (bottom) for Chr1 on simulated reads.

**Figure 20:**
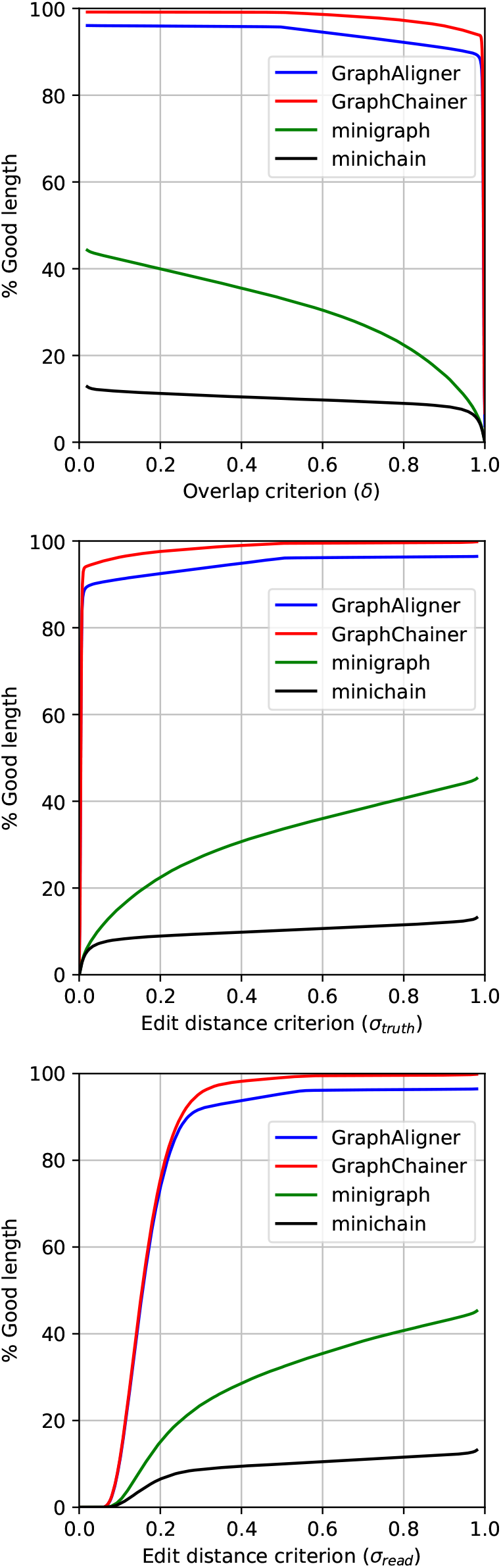
Read length in correctly aligned reads w.r.t overlap (top), truth sequence distance (middle) and read distance (bottom) for Chr1 on simulated reads.

**Figure 21:**
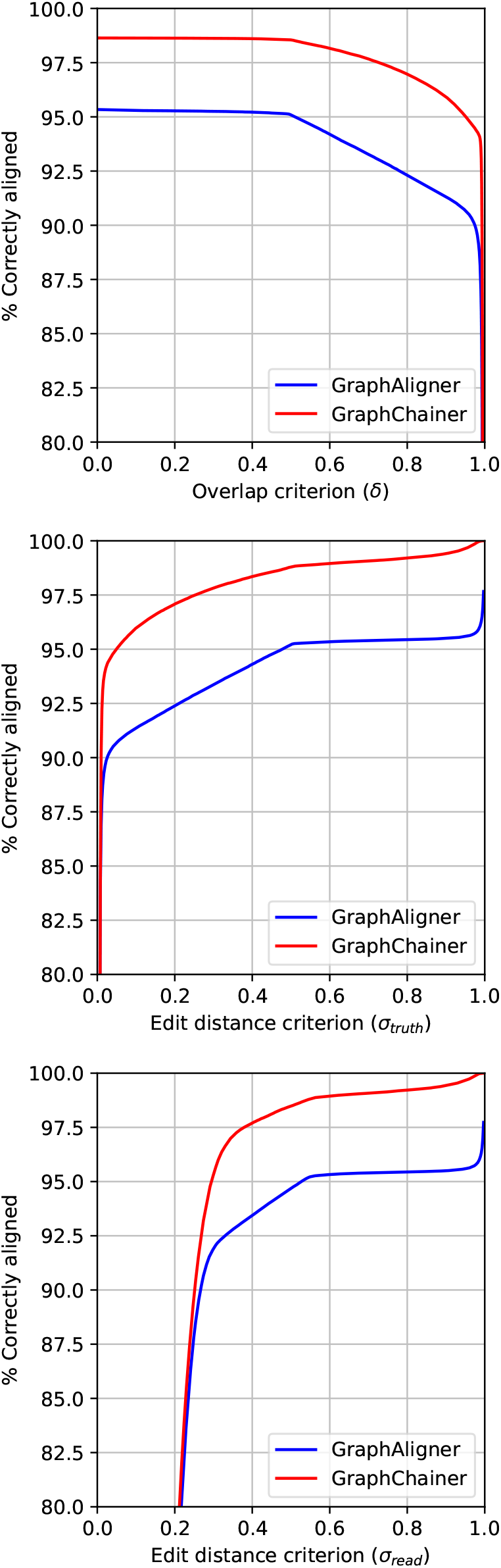
Correctly aligned reads w.r.t overlap (top), truth sequence distance (middle) and read distance (bottom) for AllChr on simulated reads (only `GraphAligner` and `GraphChainer`).

**Figure 22:**
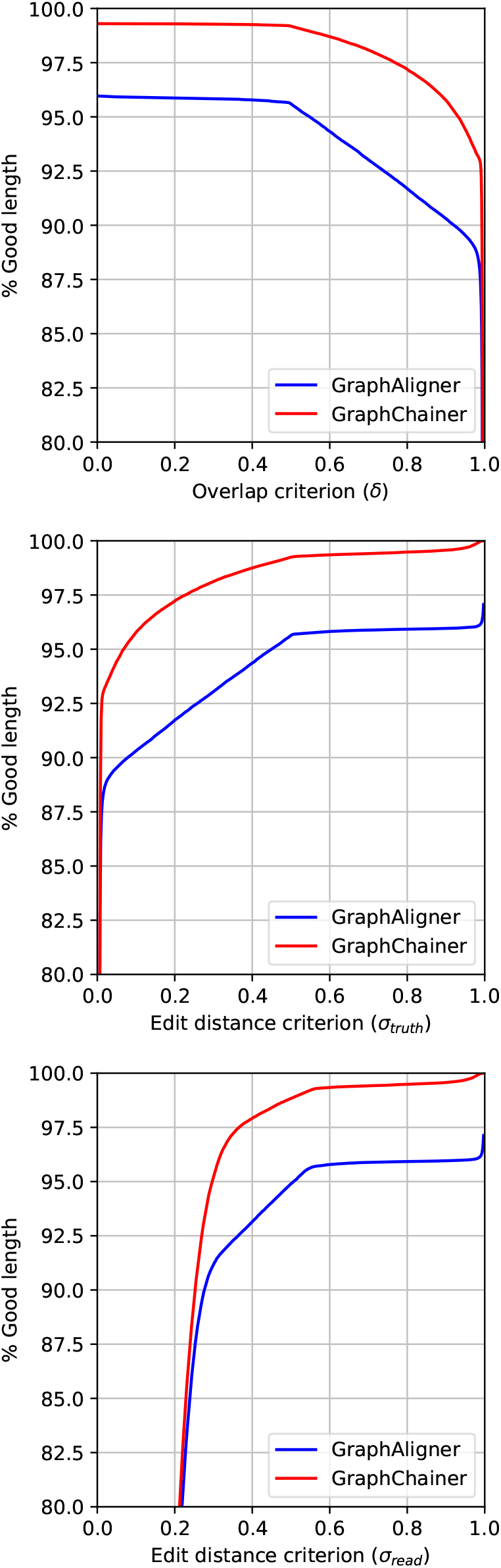
Read length in correctly aligned reads w.r.t overlap (top), truth sequence distance (middle) and read distance (bottom) for AllChr on simulated reads (only `GraphAligner` and `GraphChainer`).

**Figure 23:**
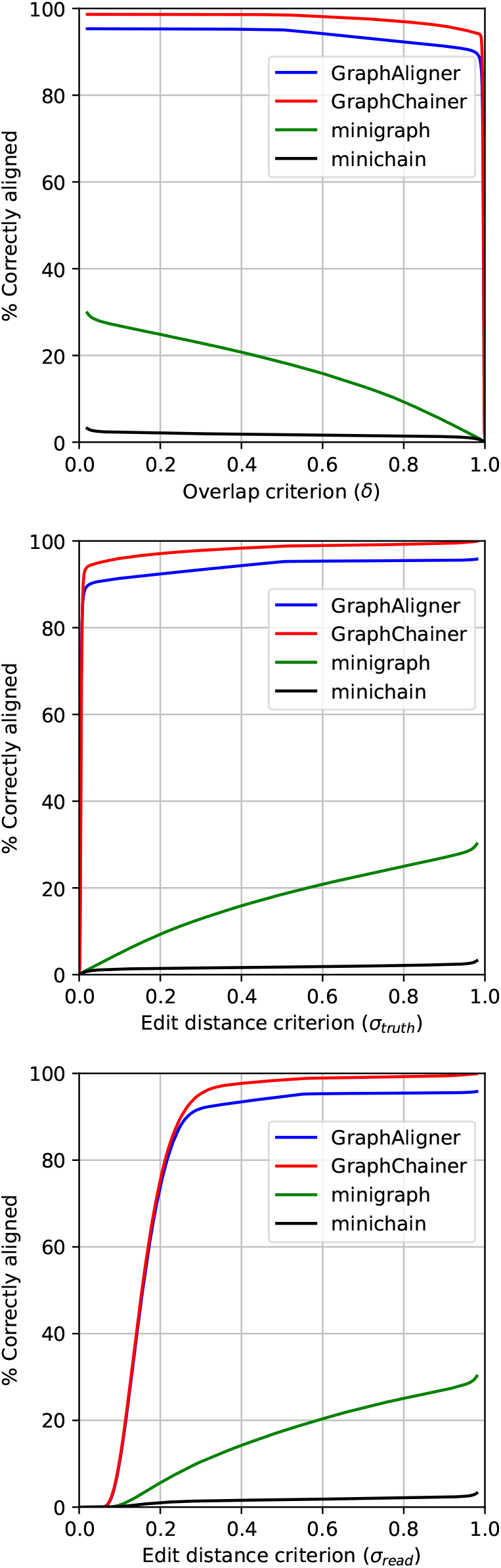
Correctly aligned reads w.r.t overlap (top), truth sequence distance (middle) and read distance (bottom) for AllChr on simulated reads.

**Figure 24:**
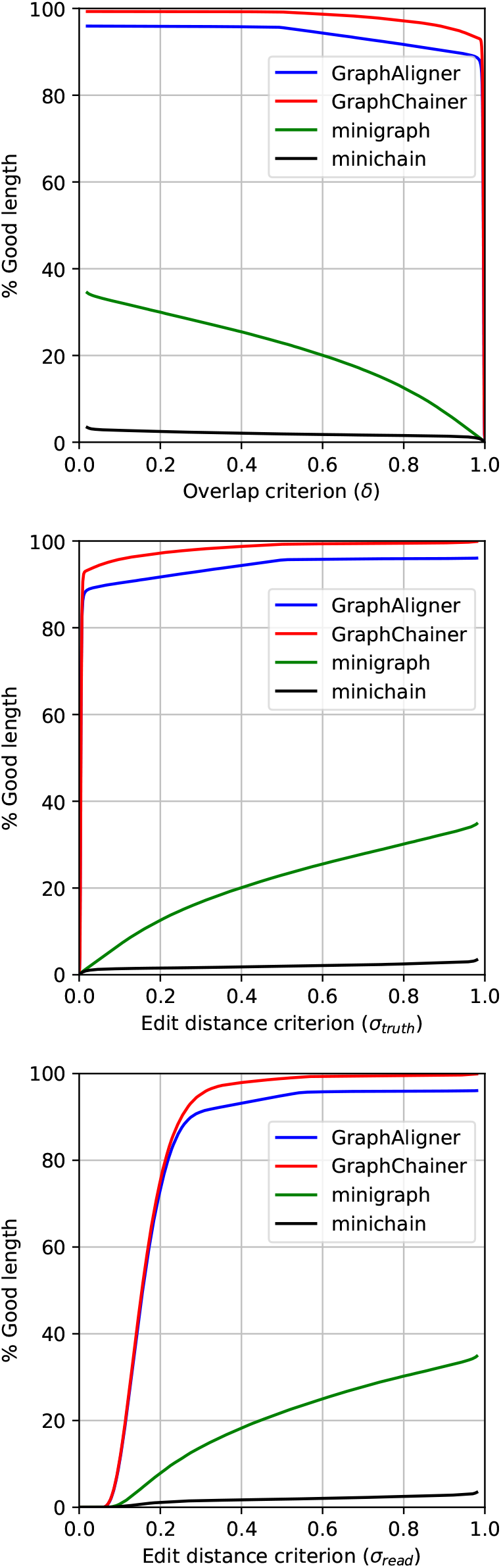
Read length in correctly aligned reads w.r.t overlap (top), truth sequence distance (middle) and read distance (bottom) for AllChr on simulated reads.

**Figure 25:**
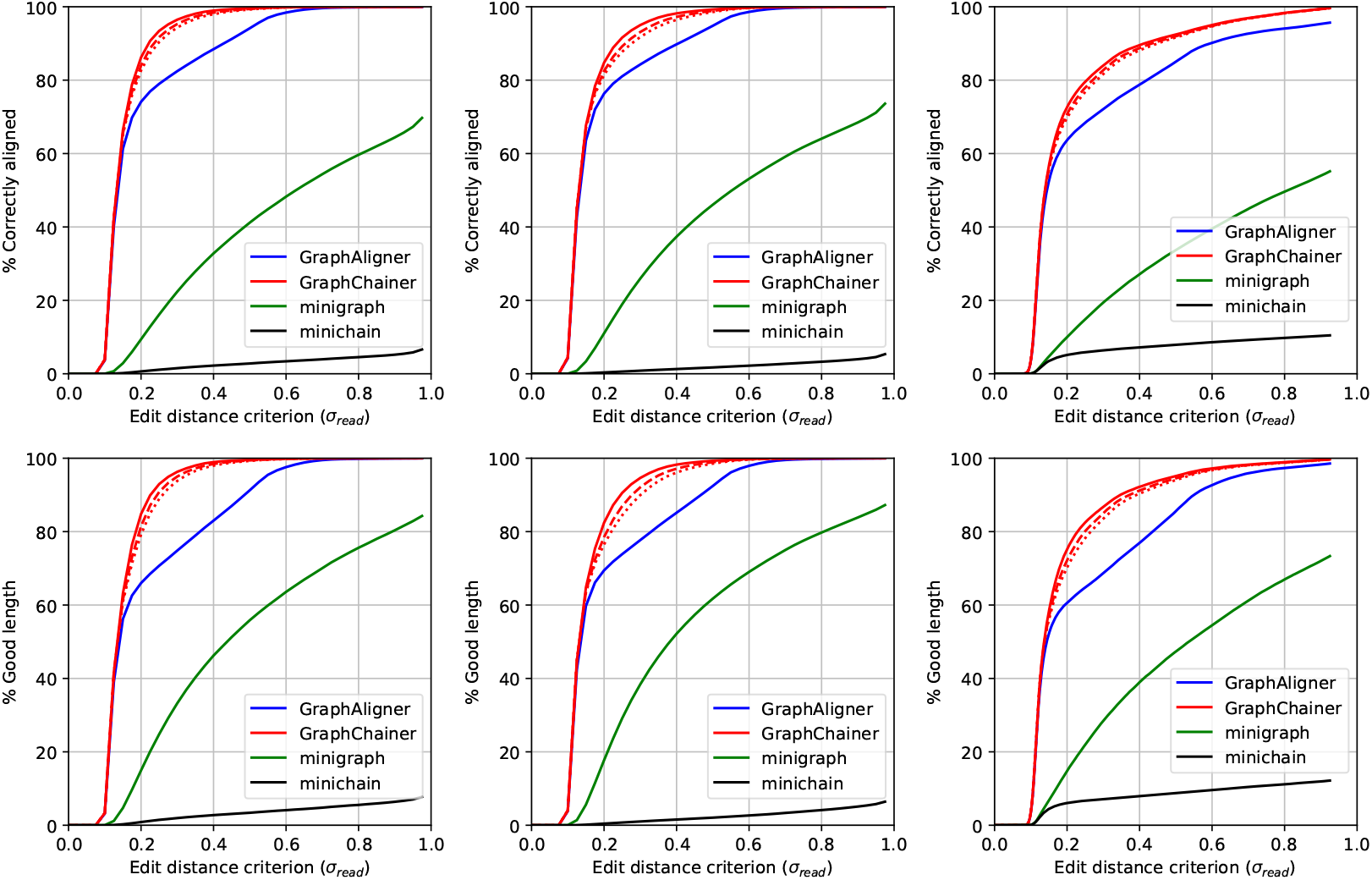
Correctly aligned reads w.r.t. the read distance (top), and read length in correctly aligned reads (bottom), on Chr22 (left), Chr1 (center) and AllChr (right), for real PacBio read sets.

**Figure 26:**
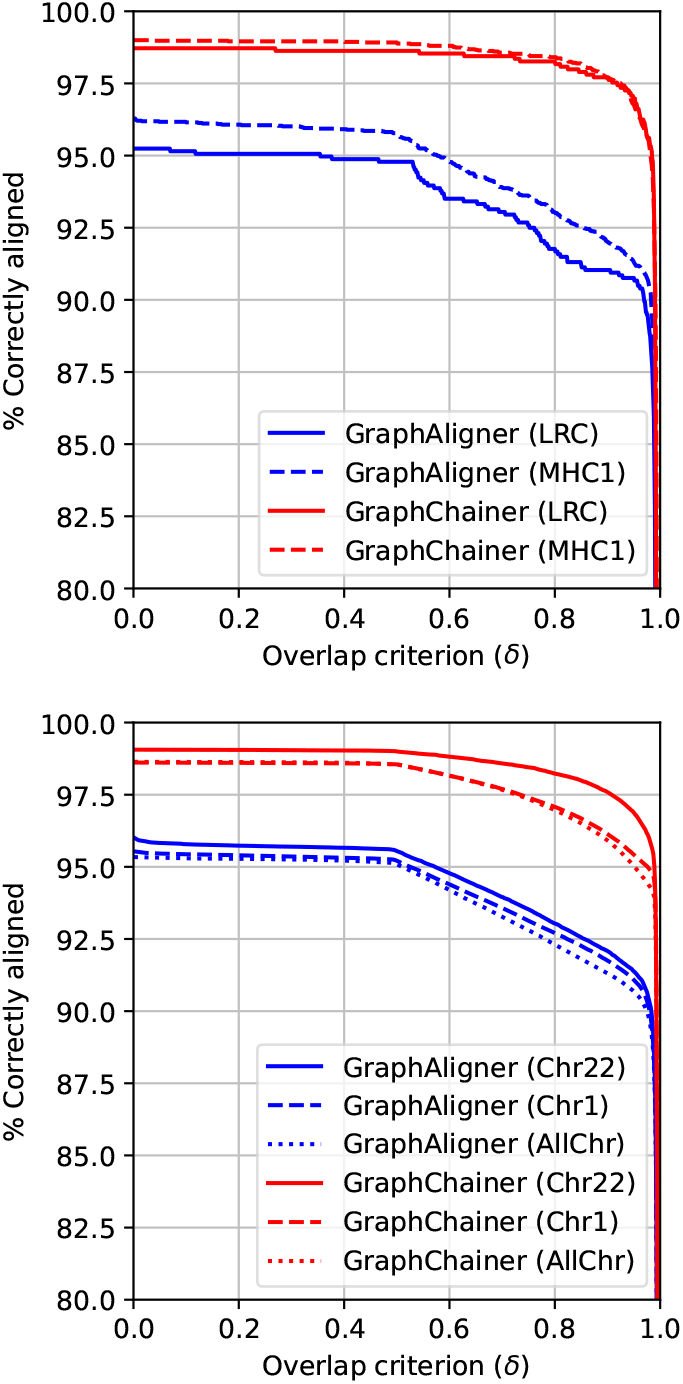
Correctly aligned reads w.r.t overlap with ground truth on the simulated read sets (5% as opposed to 15% shown in Figure 2) for LRC (top solid), MHC1 (top dashed), Chr22 (bottom solid), Chr1 (bottom dashed) and AllChr (bottom dotted).

**Figure 27:**
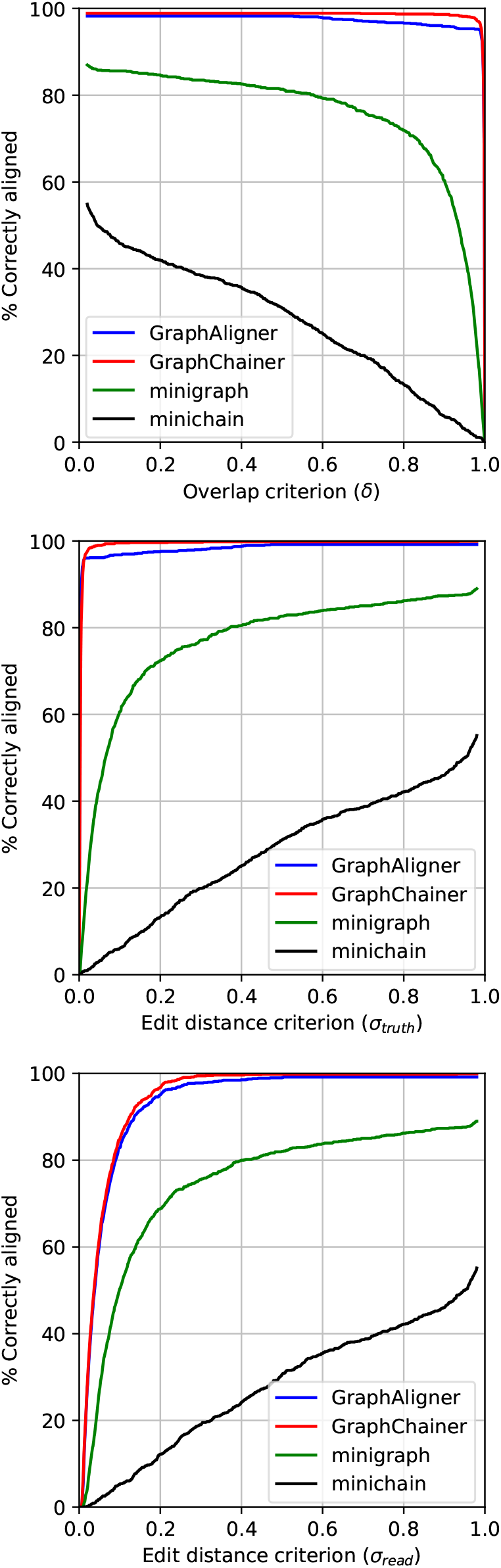
Correctly aligned reads w.r.t overlap (top), truth sequence distance (middle) (middle) and read distance (bottom) for LRC on simulated reads with error rate 5% as opposed to 15% shown in Figure 7.

**Figure 28:**
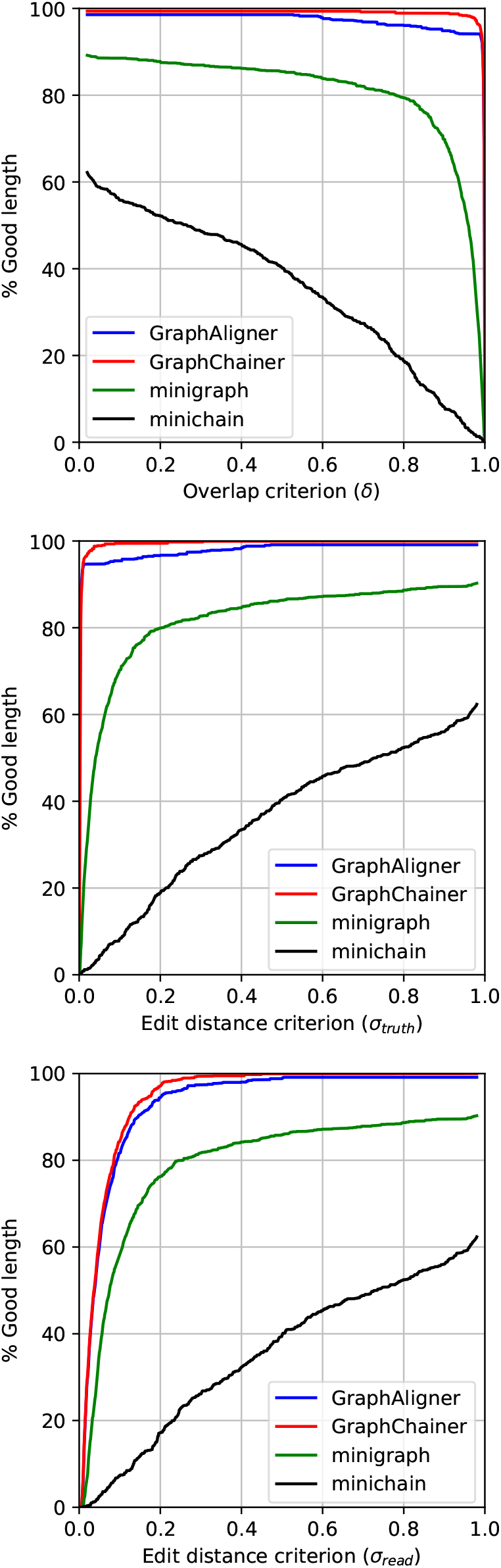
Read length in correctly aligned reads w.r.t overlap (top), truth sequence distance (middle) and read distance (bottom) for LRC on simulated reads with error rate 5% as opposed to 15% shown in Figure 8.

**Figure 29:**
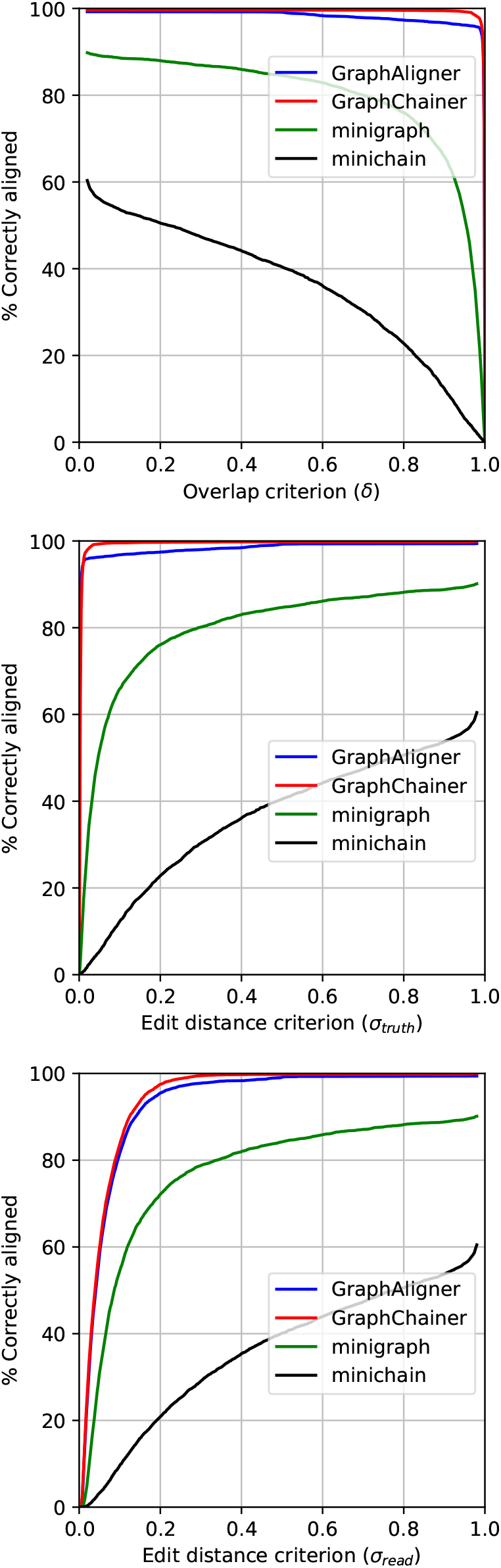
Correctly aligned reads w.r.t overlap (top), truth sequence distance (middle) and read distance (bottom) for MHC1 on simulated reads with error rate 5% as opposed to 15% shown in Figure 11.

**Figure 30:**
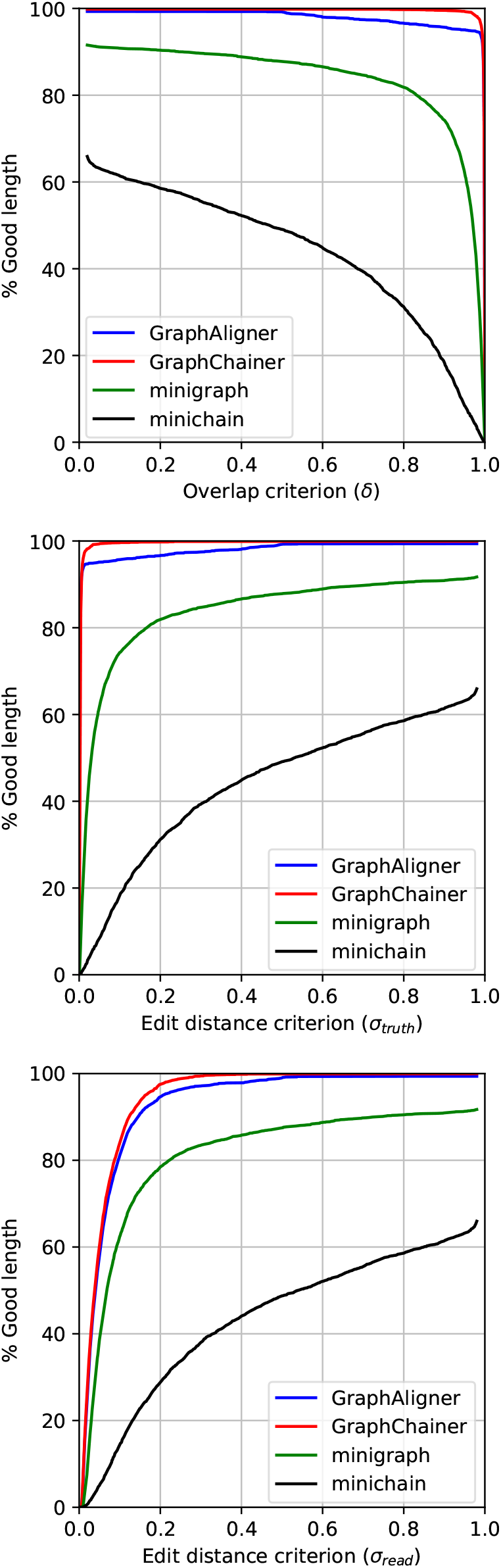
Read length in correctly aligned reads w.r.t overlap (top), truth sequence distance (middle) and read distance (bottom) for MHC1 on simulated reads with error rate 5% as opposed to 15% shown in Figure 12.

**Figure 31:**
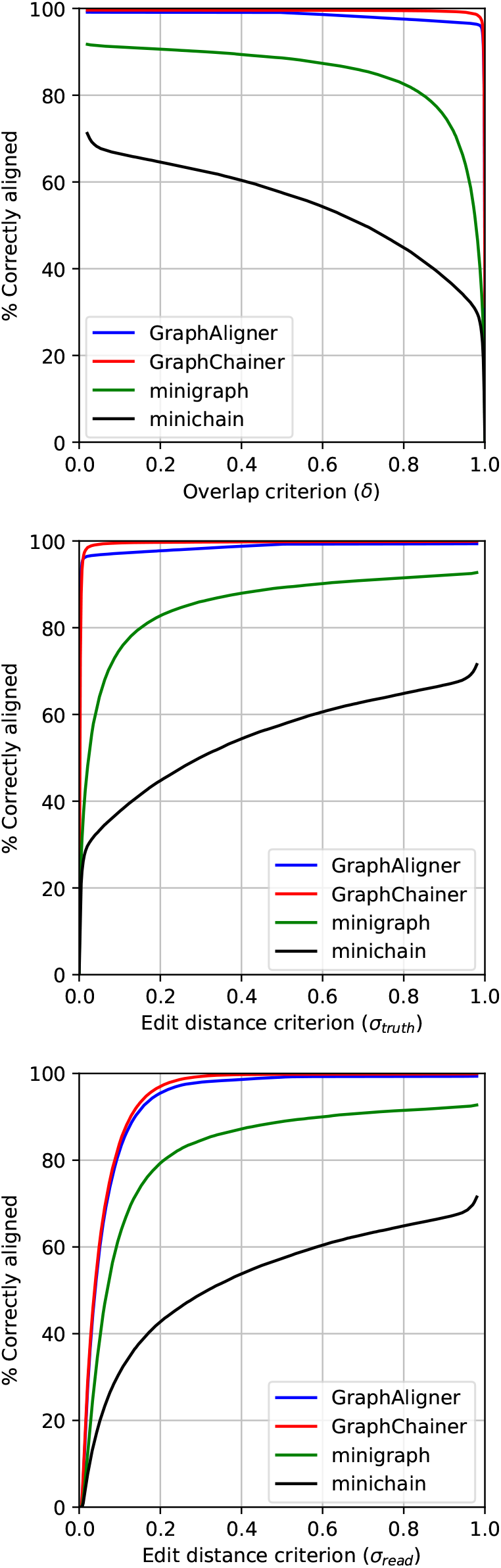
Correctly aligned reads w.r.t overlap (top), truth sequence distance (middle) and read distance (bottom) for Chr22 on simulated reads with error rate 5% as opposed to 15% shown in Figure 15.

**Figure 32:**
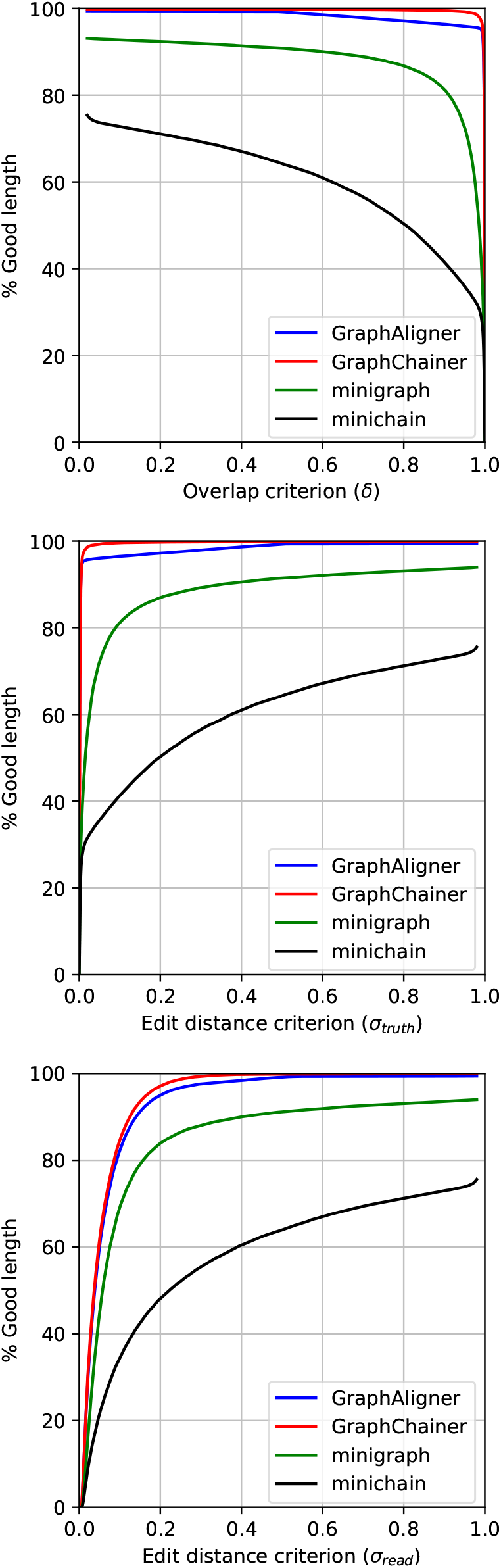
Read length in correctly aligned reads w.r.t overlap (top), truth sequence distance (middle) and read distance (bottom) for Chr22 on simulated reads with error rate 5% as opposed to 15% shown in Figure 16.

**Figure 33:**
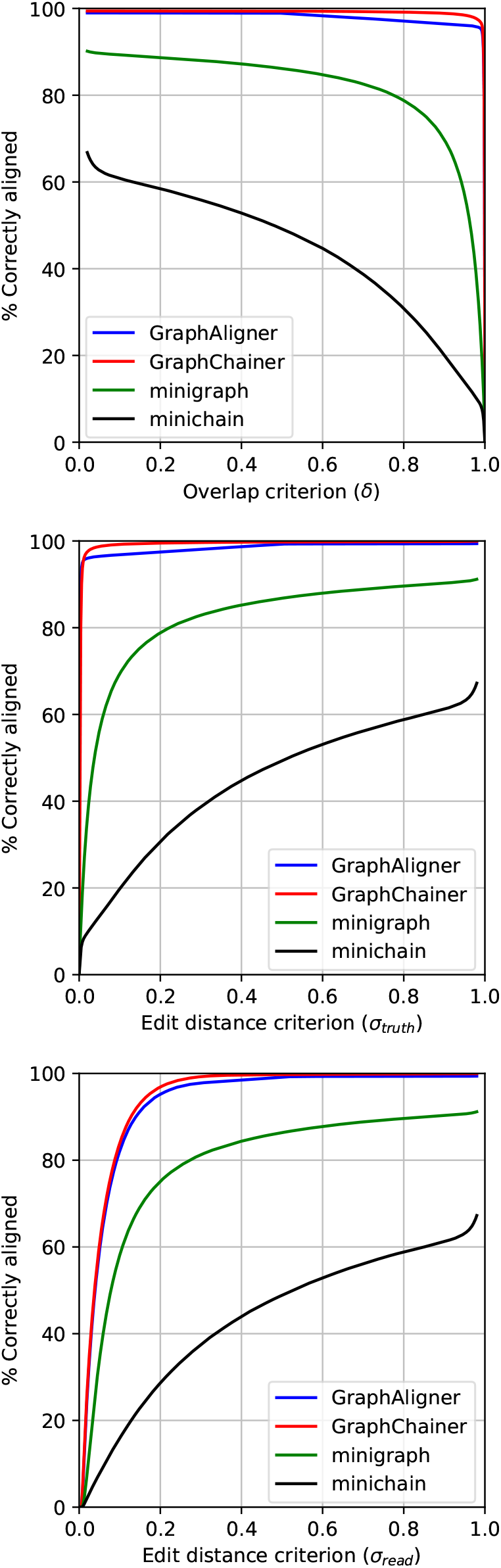
Correctly aligned reads w.r.t overlap (top), truth sequence distance (middle) and read distance (bottom) for Chr1 on simulated reads with error rate 5% as opposed to 15% shown in Figure 19.

**Figure 34:**
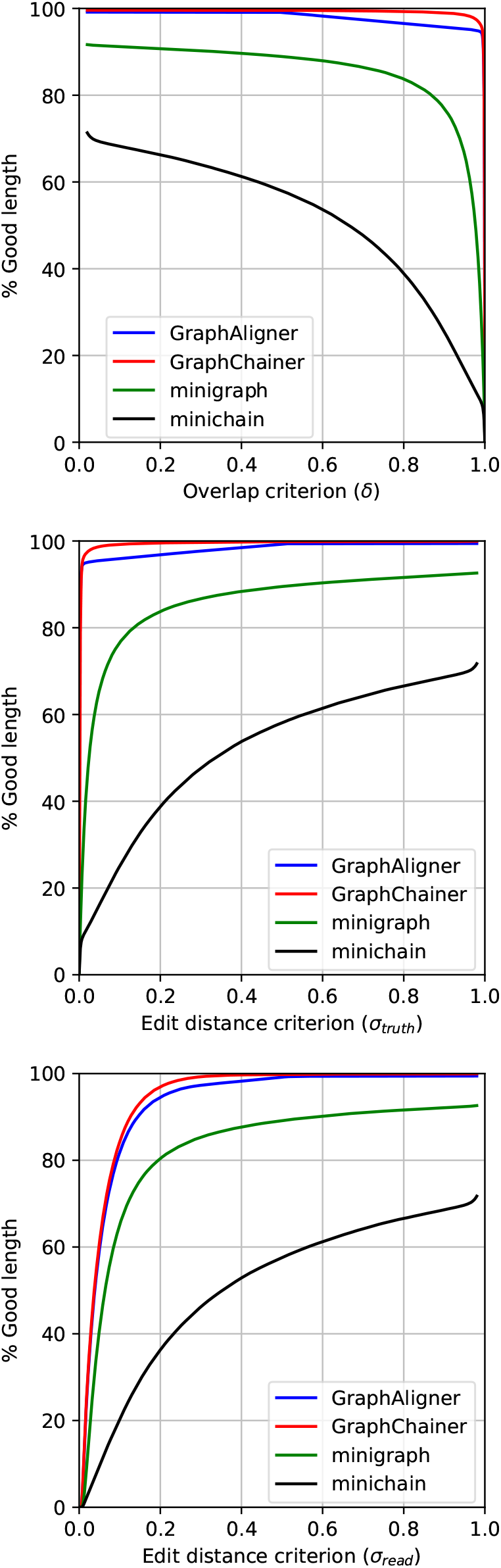
Read length in correctly aligned reads w.r.t overlap (top), truth sequence distance (middle) and read distance (bottom) for Chr1 on simulated reads with error rate 5% as opposed to 15% shown in Figure 20.

**Figure 35:**
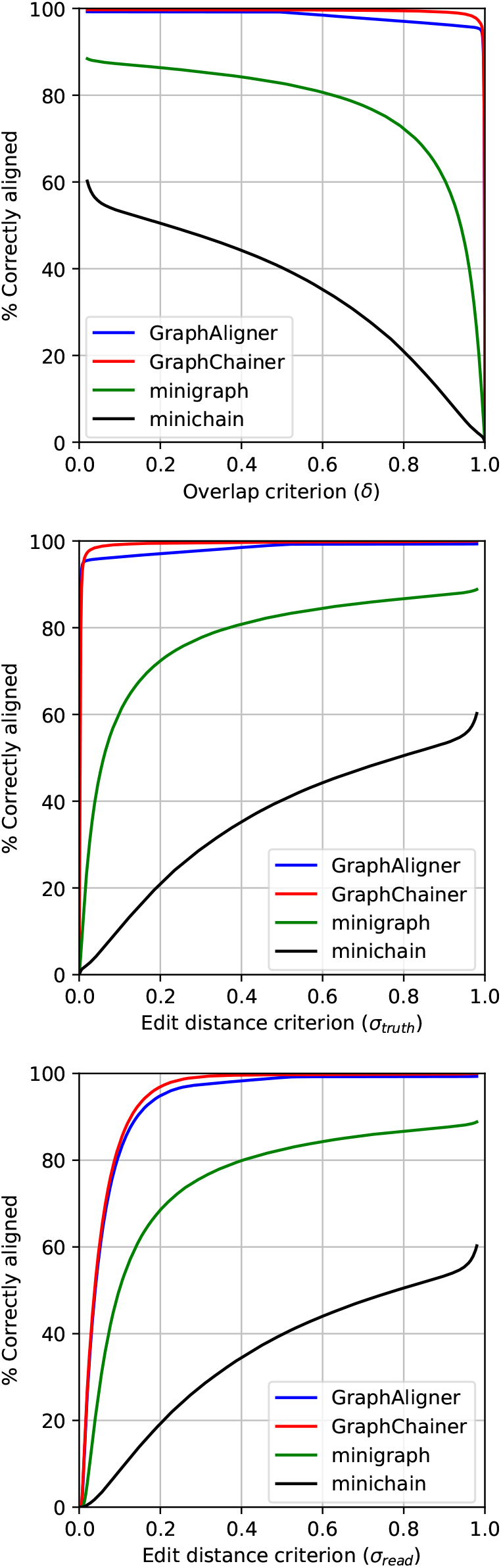
Correctly aligned reads w.r.t overlap (top), truth sequence distance (middle) and read distance (bottom) for AllChr on simulated reads with error rate 5% as opposed to 15% shown in Figure 23.

**Figure 36:**
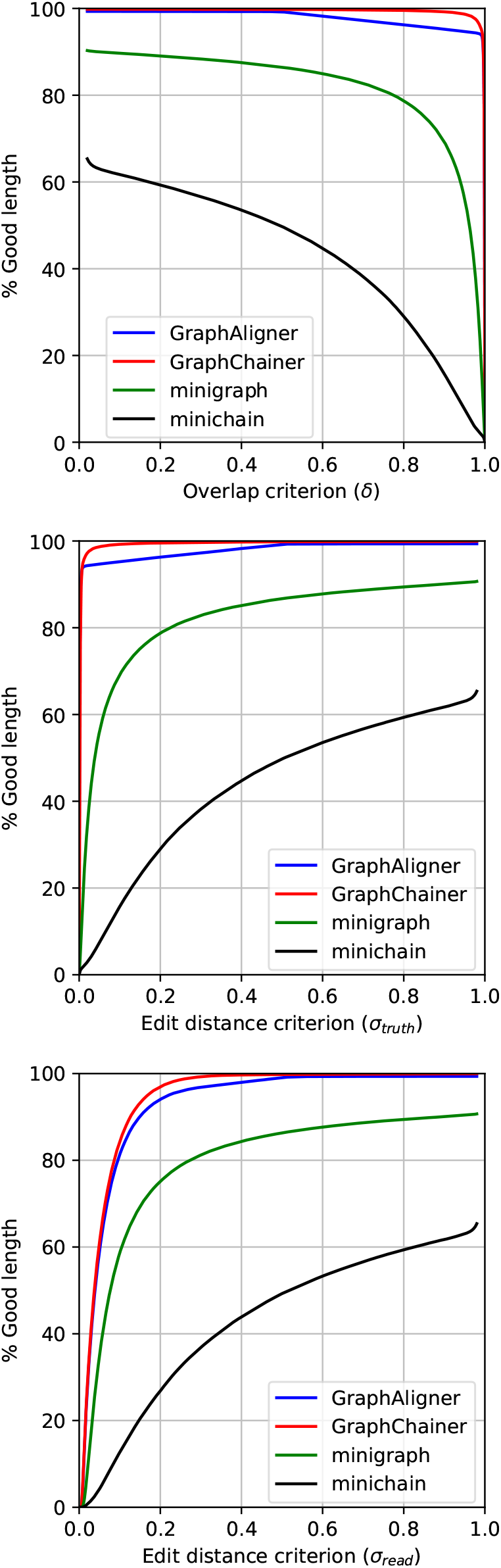
Read length in correctly aligned reads w.r.t overlap (top), truth sequence distance (middle) and read distance (bottom) for AllChr on simulated reads with error rate 5% as opposed to 15% shown in Figure 24.

**Figure 37:**
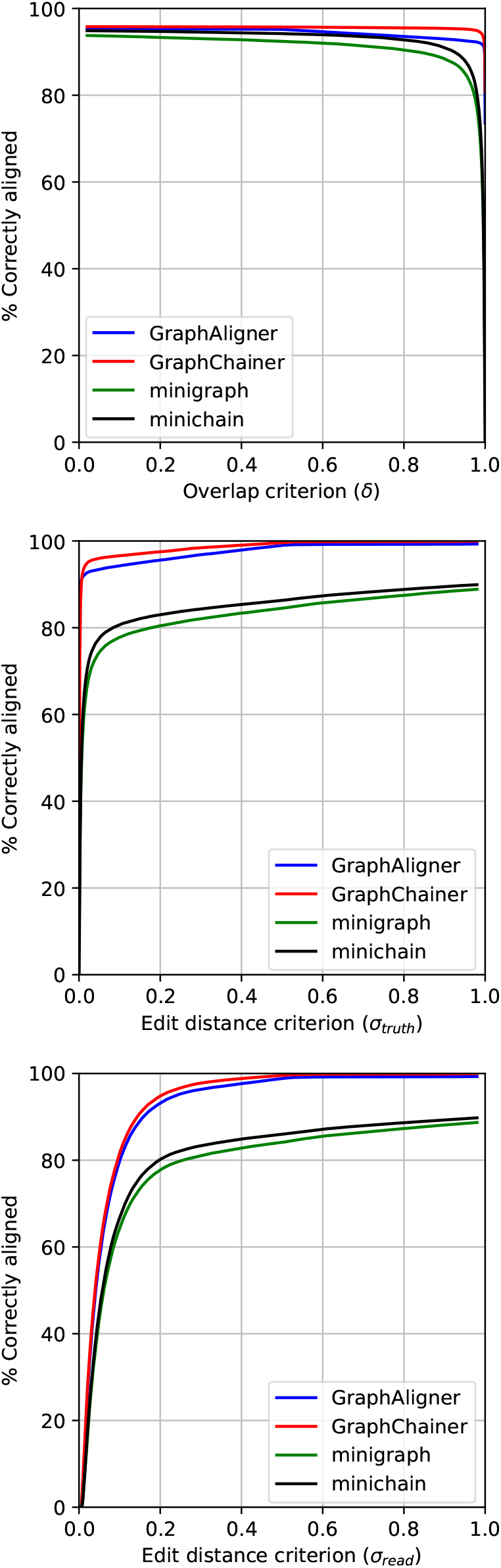
Correctly aligned reads w.r.t overlap (top), truth sequence distance (middle) and read distance (bottom) for 10H on simulated reads with error rate 5%.

**Figure 38:**
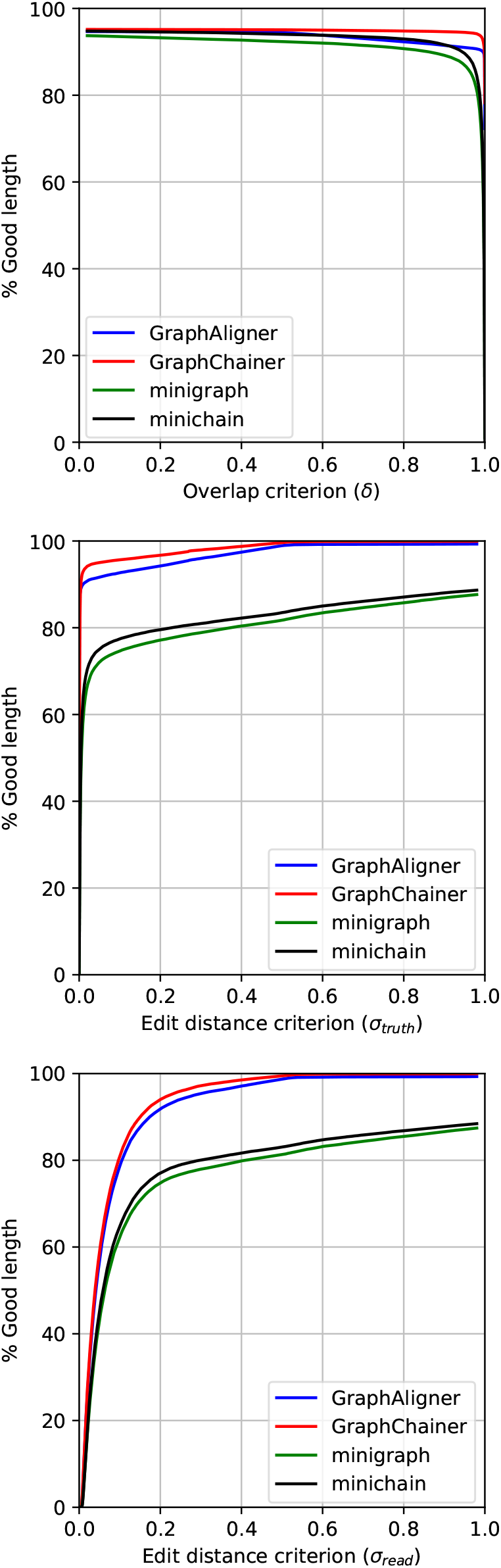
Read length in correctly aligned reads w.r.t overlap (top), truth sequence distance (middle) and read distance (bottom) for 10H on simulated reads with error rate 5%.

**Figure 39:**
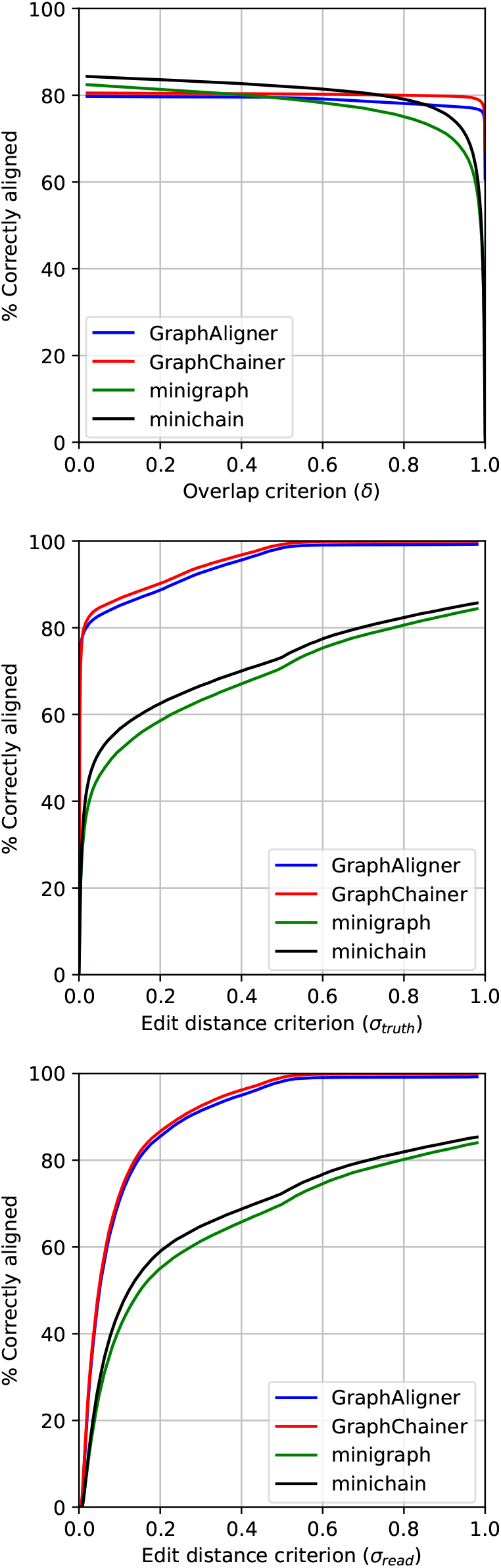
Correctly aligned reads w.r.t overlap (top), truth sequence distance (middle) and read distance (bottom) for 95H on simulated reads with error rate 5%.

**Figure 40:**
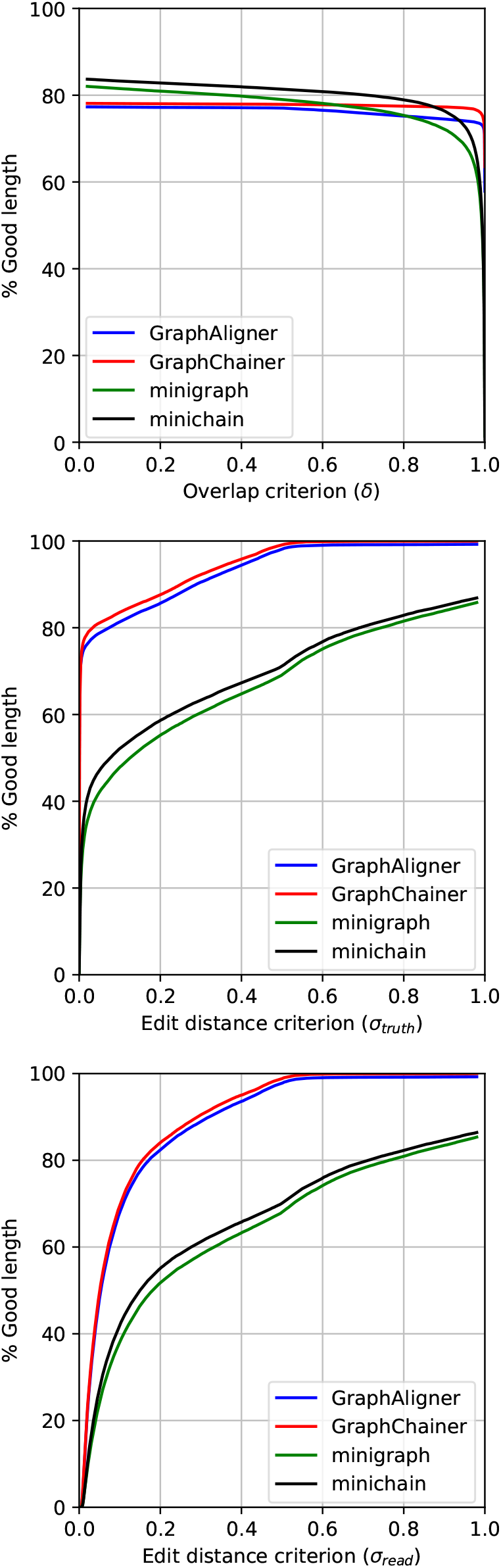
Read length in correctly aligned reads w.r.t overlap (top), truth sequence distance (middle) and read distance (bottom) for 95H on simulated reads with error rate 5%.

**Table 9:** Read length in correctly aligned reads with respect to the overlap for *δ* ∈ {0.1, 0.85} (i.e., the overlap between the reported path and the ground truth is at least 10% or 85% of the length of the ground truth sequence, respectively) for the simulated read sets. Percentages in parentheses are relative improvements w.r.t. `GraphAligner`.

| Graph | Aligner | Good length |  |
| --- | --- | --- | --- |
| | | $\delta = 0.1$ | $\delta = 0.85$ |
| LRC | **GraphAligner** | 95.67% | 90.60% |
|  | **GraphChainer** | 99.52% (+4.02%) | 98.18% (+8.37%) |
|  | minigraph | 34.26% | 8.46% |
|  | minichain | 7.51% | 0.52% |
| MHC1 | **GraphAligner** | 96.80% | 91.70% |
|  | **GraphChainer** | 99.50% (+2.79%) | 98.44% (+7.36%) |
|  | minigraph | 36.71% | 14.21% |
|  | minichain | 3.60% | 0.47% |
| Chr22 | **GraphAligner** | 96.26% | 91.91% |
|  | **GraphChainer** | 99.46% (+3.33%) | 98.19% (+6.83%) |
|  | minigraph | 53.16% | 34.32% |
|  | minichain | 32.41% | 28.17% |
| Chr1 | **GraphAligner** | 96.00% | 91.58% |
|  | **GraphChainer** | 99.17% (+3.30%) | 96.73% (+5.62%) |
|  | minigraph | 42.14% | 19.36% |
|  | minichain | 11.73% | 8.70% |
| AllChr | **GraphAligner** | 95.91% | 90.99% |
|  | **GraphChainer** | 99.30% (+3.53%) | 96.59% (+6.15%) |
|  | minigraph | 32.20% | 9.98% |
|  | minichain | 2.74% | 1.49% |

**Table 10:** Read length in correctly aligned reads with respect to the overlap for *δ* ∈ {0.1, 0.85} (i.e., the overlap between the reported path and the ground truth is at least 10% or 85% of the length of the ground truth sequence, respectively) for the simulated read sets with error rate 5% as opposed to 15% shown in Table 9. Percentages in parentheses are relative improvements w.r.t. `GraphAligner`. 10H and 95H are the smallest and largest graphs used in the experiments of `minichain`, respectively.

| Graph | Aligner | Good length |  |
| --- | --- | --- | --- |
| | | $\delta = 0.1$ | $\delta = 0.85$ |
| LRC | **GraphAligner** | 98.57% | 95.51% |
|  | **GraphChainer** | 99.37% (+0.81%) | 98.92% (+3.57%) |
|  | minigraph | 88.54% | 76.07% |
|  | minichain | 55.79% | 12.98% |
| MHC1 | **GraphAligner** | 99.30% | 96.19% |
|  | **GraphChainer** | 99.83% (+0.54%) | 99.69% (+3.64%) |
|  | minigraph | 90.80% | 78.77% |
|  | minichain | 61.38% | 25.20% |
| Chr22 | **GraphAligner** | 99.28% | 96.73% |
|  | **GraphChainer** | 99.79% (+0.51%) | 99.61% (+2.97%) |
|  | minigraph | 92.75% | 84.82% |
|  | minichain | 72.78% | 46.30% |
| Chr1 | **GraphAligner** | 99.15% | 96.14% |
|  | **GraphChainer** | 99.60% (+0.45%) | 99.20% (+3.17%) |
|  | minigraph | 91.21% | 81.29% |
|  | minichain | 68.23% | 32.94% |
| AllChr | **GraphAligner** | 99.33% | 95.77% |
|  | **GraphChainer** | 99.81% (+0.49%) | 99.44% (+3.84%) |
|  | minigraph | 89.68% | 75.16% |
|  | minichain | 61.71% | 22.91% |
| 10H | **GraphAligner** | 94.64% | 91.94% |
|  | **GraphChainer** | 95.15% (+0.54%) | 94.75% (+3.06%) |
|  | minigraph | 93.52% | 90.18% |
|  | minichain | 94.71% | 92.52% |
| 95H | **GraphAligner** | 77.26% | 74.87% |
|  | **GraphChainer** | 78.08% (+1.06%) | 77.38% (+3.36%) |
|  | minigraph | 81.52% | 74.08% |
|  | minichain | 83.26% | 78.01% |

**Table 11:**
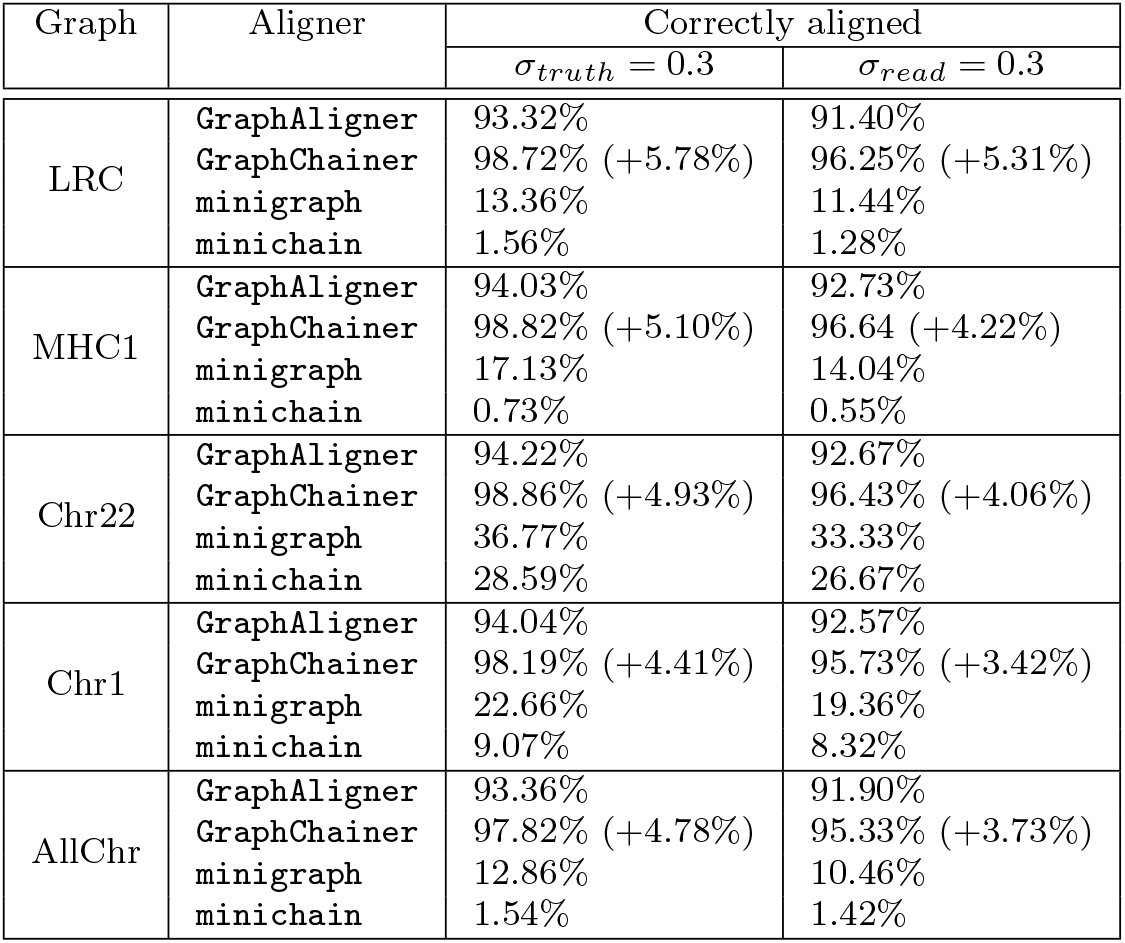
Correctly aligned reads with respect to the distance, for *σ*_*truth*_ = 0.3 (i.e., the edit distance between the truth sequence and the reported sequence can be up to 30% of the truth sequence length), and *σ*_*read*_ = 0.3 (i.e., the edit distance between the read and the reported sequence can be up to 30% of the read length) for simulated read sets. Percentages in parentheses are relative improvements w.r.t. `GraphAligner`.

| Graph | Aligner | Correctly aligned |  |
| --- | --- | --- | --- |
| | | $\sigma_{truth} = 0.3$ | $\sigma_{read} = 0.3$ |
| LRC | **GraphAligner** | 93.32% | 91.40% |
|  | **GraphChainer** | 98.72% (+5.78%) | 96.25% (+5.31%) |
|  | minigraph | 13.36% | 11.44% |
|  | minichain | 1.56% | 1.28% |
| MHC1 | **GraphAligner** | 94.03% | 92.73% |
|  | **GraphChainer** | 98.82% (+5.10%) | 96.64 (+4.22%) |
|  | minigraph | 17.13% | 14.04% |
|  | minichain | 0.73% | 0.55% |
| Chr22 | **GraphAligner** | 94.22% | 92.67% |
|  | **GraphChainer** | 98.86% (+4.93%) | 96.43% (+4.06%) |
|  | minigraph | 36.77% | 33.33% |
|  | minichain | 28.59% | 26.67% |
| Chr1 | **GraphAligner** | 94.04% | 92.57% |
|  | **GraphChainer** | 98.19% (+4.41%) | 95.73% (+3.42%) |
|  | minigraph | 22.66% | 19.36% |
|  | minichain | 9.07% | 8.32% |
| AllChr | **GraphAligner** | 93.36% | 91.90% |
|  | **GraphChainer** | 97.82% (+4.78%) | 95.33% (+3.73%) |
|  | minigraph | 12.86% | 10.46% |
|  | minichain | 1.54% | 1.42% |

**Table 12:** Correctly aligned reads with respect to the distance, for *σ*_*truth*_ = 0.3 (i.e., the edit distance between the truth sequence and the reported sequence can be up to 30% of the truth sequence length), and *σ*_*read*_ = 0.3 (i.e., the edit distance between the read and the reported sequence can be up to 30% of the read length) for simulated read sets with error rate 5% as opposed to 15% shown in Table 11. Percentages in parentheses are relative improvements w.r.t. `GraphAligner`. 10H and 95H are the smallest and largest graphs used in the experiments of `minichain`, respectively.

| Graph | Aligner | Correctly aligned |  |
| --- | --- | --- | --- |
| | | $\sigma_{truth} = 0.3$ | $\sigma_{read} = 0.3$ |
| LRC | **GraphAligner** | 97.98% | 97.80% |
|  | **GraphChainer** | 99.72% (+1.78%) | 99.45% (+1.69%) |
|  | **minigraph** | 77.16% | 75.50% |
|  | **minichain** | 19.91% | 18.99% |
| MHC1 | **GraphAligner** | 98.02% | 97.74% |
|  | **GraphChainer** | 99.76% (+1.78%) | 99.49% (+1.78%) |
|  | **minigraph** | 80.16% | 78.81% |
|  | **minichain** | 30.30% | 29.07% |
| Chr22 | **GraphAligner** | 98.26% | 97.99% |
|  | **GraphChainer** | 99.74% (+1.51%) | 99.31% (+1.34%) |
|  | **minigraph** | 86.01% | 84.57% |
|  | **minichain** | 50.16% | 49.13% |
| Chr1 | **GraphAligner** | 98.09% | 97.79% |
|  | **GraphChainer** | 99.64% (+1.58%) | 99.27% (+1.50%) |
|  | **minigraph** | 82.87% | 81.25% |
|  | **minichain** | 38.73% | 37.49% |
| AllChr | **GraphAligner** | 97.76% | 97.46% |
|  | **GraphChainer** | 99.61% (+1.89%) | 99.24% (+1.83%) |
|  | **minigraph** | 77.71% | 75.86% |
|  | **minichain** | 28.98% | 27.85% |
| 10H | **GraphAligner** | 96.82% | 96.37% |
|  | **GraphChainer** | 98.42% (+1.65%) | 97.78% (+1.47%) |
|  | **minigraph** | 82.06% | 81.08% |
|  | **minichain** | 84.32% | 83.35% |
| 95H | **GraphAligner** | 92.67% | 91.42% |
|  | **GraphChainer** | 94.06% (+1.50%) | 92.54% (+1.22%) |
|  | **minigraph** | 63.27% | 61.35% |
|  | **minichain** | 66.67% | 64.79% |

**Table 13:** Read length in correctly aligned reads with respect to the distance, for *σ*_*truth*_ = 0.3 (i.e., the edit distance between the truth sequence and the reported sequence can be up to 30% of the truth sequence length), and *σ*_*read*_ = 0.3 (i.e., the edit distance between the read and the reported sequence can be up to 30% of the read length) for simulated read sets. Percentages in parentheses are relative improvements w.r.t. `GraphAligner`.

| Graph | Aligner | Good length |  |
| --- | --- | --- | --- |
| | | $\sigma_{truth} = 0.3$ | $\sigma_{read} = 0.3$ |
| LRC | **GraphAligner** | 92.79% | 90.47% |
|  | **GraphChainer** | 99.08% (+6.78%) | 95.87% (+5.97%) |
|  | minigraph | 17.09% | 14.61% |
|  | minichain | 1.38% | 1.05% |
| MHC1 | **GraphAligner** | 93.42% | 92.06% |
|  | **GraphChainer** | 99.00% (+5.97%) | 96.76% (+5.10%) |
|  | minigraph | 21.88% | 18.62% |
|  | minichain | 0.92% | 0.70% |
| Chr22 | **GraphAligner** | 93.97% | 92.03% |
|  | **GraphChainer** | 99.05% (+5.41%) | 96.35% (+4.69%) |
|  | minigraph | 40.49% | 37.04% |
|  | minichain | 29.55% | 27.95% |
| Chr1 | **GraphAligner** | 93.72% | 91.87% |
|  | **GraphChainer** | 98.39% (+4.99%) | 95.58% (+4.03%) |
|  | minigraph | 27.20% | 23.58% |
|  | minichain | 9.39% | 8.72% |
| AllChr | **GraphAligner** | 93.04% | 91.17% |
|  | **GraphChainer** | 98.13% (+5.47%) | 95.22% (+4.42%) |
|  | minigraph | 16.72% | 13.87% |
|  | minichain | 1.65% | 1.54% |

**Table 14:**
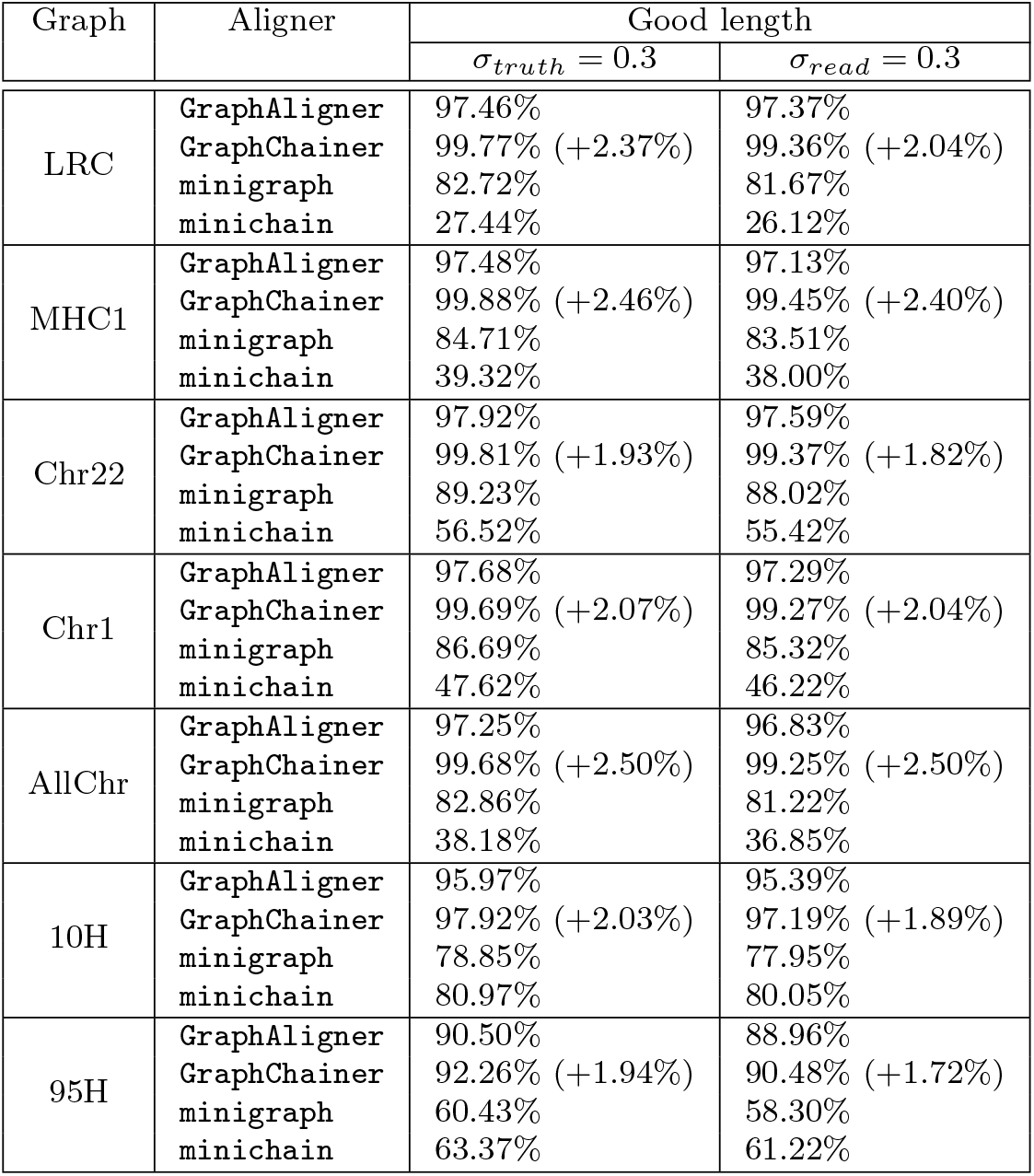
Read length in correctly aligned reads with respect to the distance, for *σ*_*truth*_ = 0.3 (i.e., the edit distance between the truth sequence and the reported sequence can be up to 30% of the truth sequence length), and *σ*_*read*_ = 0.3 (i.e., the edit distance between the read and the reported sequence can be up to 30% of the read length) for simulated read sets with error rate 5% as opposed to 15% shown in Table 13. Percentages in parentheses are relative improvements w.r.t. `GraphAligner`. 10H and 95H are the smallest and largest graphs used in the experiments of `minichain`, respectively.

## Footnotes

1 Highly variable whole human genome graphs of thousands of individuals.

2 *L* corresponds to the sum of the path lengths, whereas #*o* is the number of overlaps between paths.

3 Although our graphs have high variability, it is worth noting that they are acyclic excluding structural variants such as repeat expansions, inversions, long deletions and insertions.

4 Two paths have a suffix-prefix overlap if the last *k* vertices of one (suffix) are exactly the last *k* vertices of the other (prefix), for some *k >* 0.

5 *L* corresponds to the sum of the path lengths, whereas #*o* is the number of overlaps between paths.

6 Two paths have a one-node suffix-prefix overlap if the last vertex of one is exactly the first vertex of the other.

7 We refer to the original publication [33] for the exact information stored and queried in the data structure, or to our pseudocode in Algorithm 1.

8 We refer to the original publication [33] for the exact definition and construction algorithm for the *forward propagation links*, or to our pseudocode in Algorithm 1.

9 All calls to `GraphAligner` are implemented internally, in the same binary of `GraphChainer`.

10 We ignore the alignment score returned by `GraphAligner`. Other aligners such as `minigraph` and `minichain` consider this score as part of the optimization criterion of co-linear chaining.

11 Only 5% error rate for the case of 10H and 95H to replicate `minichain`’s experimental setting.

12 This is the same read set used by `GraphAligner`’s experiments [37, p.7] but without the subsampling to 15 *×* coverage.

13 Both objects, as well as the ground truth path/sequence, exclude the offsets of the first and last nodes.

14 Also on simulated data to show the relation between the criteria. This can be found in the Appendix.

15 Recall that real reads used for AllChr are a uniform random sample of real reads used for Chr1.

16 For presentation reasons we keep these results in the tables but exclude them from the plots. The corresponding plots can be found in the Appendix (see Figures 7, 8, 11, 12, 15, 16, 19, 20, 23 and 24).

17 These can be found in the Appendix: Tables 5, 10, 12 and 14 and Figures 26 to 40.

18 Note that we worsen running time by a *k* factor inside the log factor. However, *k* ≪ *N*, thus the asymptotic running time is maintained.

19 If there are ties, they are sorted by endpoint in the string label of the path endpoint. This is done in this case as well as the other sortings, but we remove it from the phrasing by ease of explanation.

